# Mechanistic insights into coronavirus polymerase inhibition by a base-modified adenosine analog

**DOI:** 10.64898/2026.10.28.761875

**Authors:** Nikolai Bakirtzoglou, Barbara Selisko, Yucen Huang, Peng Nie, Bruno Ribeiro de Castro Ramos, Ria Van Berwaer, Lieselotte Schurmans, Veronique Fattorini, Yu Zhou, Yufan Pan, Xiaoxia An, Julie Vandeput, Benjamin Van Loy, Bert Vanmechelen, Piet Maes, Dongsheng Liu, Piet Herdewijn, Bruno Canard, Zhiyong Lou, Elisabetta Groaz, Lieve Naesens, Annelies Stevaert

## Abstract

Nucleoside analog (NA) inhibitors of viral polymerases are a chemically diverse and prominent class of antivirals, with most agents bearing sugar modifications to induce chain termination. Compounds with a modified base yet natural ribose are comparatively rare and typically incorporated to act as virus mutagens, raising significant safety concerns. How nucleobase modifications interfere with polymerization and the molecular basis of this effect remain poorly understood. We here define the mechanism of 6-ethyl-modified 7-deazaadenosine (EDA), a recently identified coronavirus (CoV) inhibitor that does not induce virus mutagenesis. Its strong activity spans across multiple CoVs and cell models and is dictated by the 6-ethyl substituent, with poor tolerance for smaller, larger or heteroatom-containing substituents. At the level of SARS-CoV-2 RNA-dependent RNA polymerase (RdRp), EDA-triphosphate incorporates as a mimic of ATP but not other NTPs, inducing immediate stalling but also excision by the viral 3′-to-5′ exonuclease. While EDA incorporation prevents extension with a second EDA residue, the stalling of natural NTP incorporation is transient and still allows the formation of longer products. A similar phenomenon is observed with template-embedded EDA. Cryo-EM analysis with SARS-CoV-2 RdRp revealed that 3′-terminal EDA is accommodated without distortion of the template:primer duplex, but its 6-ethyl group induces rearrangements in the NTP binding pocket that prevent stable binding of the incoming nucleotide. The mechanistic findings were validated through antiviral studies with SARS-CoV-2 mutants bearing core substitutions in the RdRp or exonuclease. Together, these findings position EDA as a mechanistically distinct base-modified NA prototype that has relevance beyond coronaviruses and underscores the ever-expanding versatility of NA-based polymerase inhibitors.

## Introduction

Despite broad availability of vaccines and the establishment of population-level immunity, COVID-19 remains a significant clinical challenge, particularly for fragile persons such as those of older age or suffering from comorbidities [1]. In these settings, the disease burden of COVID-19 is still at least as high as that of seasonal influenza [2]. The first-line oral therapy, nirmatrelvir boosted with ritonavir, may lead to viral resistance [3] or post-treatment viral rebound [4]. The alternative drugs, approved in certain countries, comprise the nucleoside analogs (NAs) remdesivir (RDV), favipiravir and molnupiravir. The latter two have ambiguous base-pairing properties, explaining why their incorporation into coronavirus genomic RNA results in (lethal) virus mutagenesis [5–7]. Since they might also induce mutations in cellular RNA or DNA, their clinical use is not free of safety concerns [8]. Besides, broad use of mutagenic NAs may alter viral evolution in the human population [9]. Consequently, there remains a strong need for NAs with strong anti-coronavirus (CoV) activity and a safe clinical profile.

The clinical success of antiviral NAs can be attributed to their broad applicability and high versatility in chemical structure [10] and mechanism of action [11–13]. Most of them act as inhibitors of viral polymerases but the precise mechanism shows considerable variation, depending also on assay conditions. RDV was shown to cause ‘delayed’ stalling of the SARS-CoV-2 RNA-dependent RNA polymerase [RdRp; non-structural protein 12 (nsp12)] [14–16], due to a steric clash between the enzyme and 1′-cyano function of RDV [14, 17]. Still, under physiological NTP concentrations, the RdRp can surmount the stalling by RDV [18], which likely results in RDV integration into viral (sub)genomic RNA and impairment of next-round RNA synthesis when the RdRp encounters a templating RDV [19]. Second, the CoV RdRp functions within a protein complex that includes the viral 3′-to-5′ exonuclease (ExoN; nsp14) proofreading enzyme [20], which excises mismatched nucleotides [21] and is critical for faithful replication of the large CoV RNA genome [22]. The ExoN activity complicates the development of NAs, since unnatural substitutions in the ribose or base part may trigger fast excision of the compound, reducing its antiviral potency [23]. Whereas RDV encounters some though limited impact from ExoN removal [24], molnupiravir appears hardly affected [7]. The excision rate of an NA is likely dictated by a combination of its chemical structure and inhibitory mechanism at RdRp level [25], but remains challenging to predict despite its importance for developing highly active anti-CoV NAs.

In an effort to develop original NA scaffolds with interesting pharmacological properties, a series of 6-alkyl- and 6-aryl-substituted 7-deazapurine ribonucleoside compounds was synthesized a few years ago [26] and, along with a series of 6-alkynyl substituted analogs, evaluated against a broad range of RNA viruses [27, 28]. One compound that emerged as particularly appealing is the 6-ethyl analog, consisting of 7-deazaadenosine in which the exocyclic NH_2_ function is replaced by an ethyl moiety. This molecule, abbreviated as EDA (Fig 1A), was shown to produce robust inhibition of influenza A and B viruses, with antiviral EC_50_ values in the submicromolar range [27, 28]. The compound proved also effective in a replicon assay with SARS-CoV-2, indicating an appealing profile of dual activity against influenza and CoVs [28].

**Fig 1.**
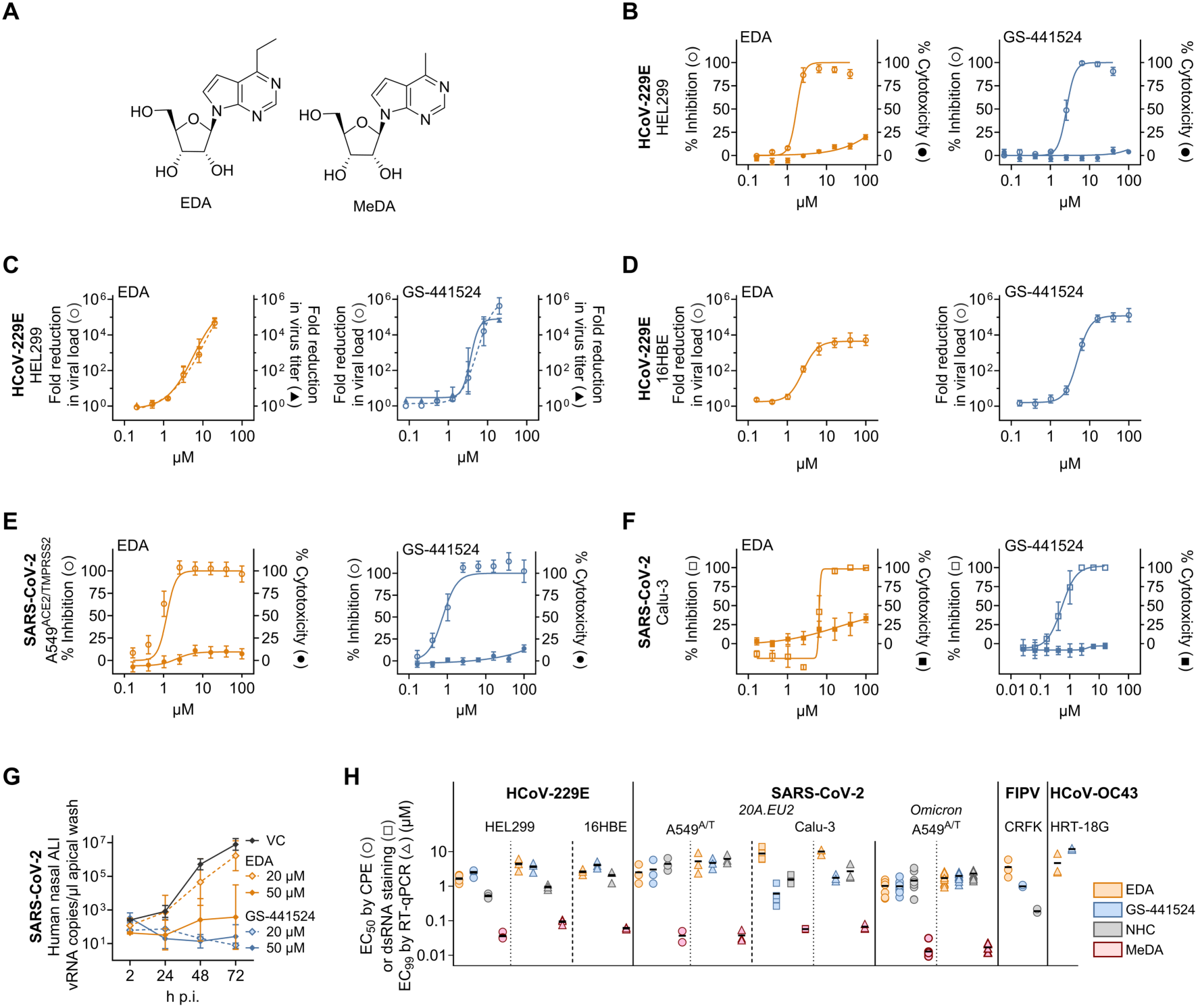
EDA has broad activity across multiple CoVs and cell models. (**A**) Chemical structures of EDA and MeDA. (**B**-**C**) Inhibition of HCoV-229E and cytotoxicity in HEL299 cells, based on cell viability assay (panel **B**), or assessment of viral load and infectious titer (panel **C**). (**D**) Anti-HCoV-229E activity in human bronchial 16HBE cells. (**E**-**G**) Inhibition of SARS-CoV-2 and cytotoxicity, based on cell viability assay in A549^ACE2/TMPRSS2^ (A549A/T) cells (panel **E**); immunostaining for viral dsRNA in human bronchial Calu-3 cells (panel **F**); or determination of viral load in primary human nasal cells grown at air-liquid interface (panel **G**). Data in panels B-G are mean values ± SEM (N=3-5 except for: panel E, where N=10; and panel G, where N=2-3). (**H**) Compilation of the anti-CoV activity values of EDA, MeDA, GS-441524 and NHC, across different CoVs and cell models. Symbols: EC_50_ or EC_99_ values, depending on the assay readout, from 2-13 independent tests; black bar: mean value.

Since EDA possesses a natural ribose, it does not qualify as an ‘obligate’ chain terminator, a term used for NAs that lack a 3′-hydroxyl group [29]. It also differs from known ‘non-obligate’ chain terminators which, typically, carry modifications at other positions of the ribose (e.g., 1′, 2′ or 4′). Antiviral NAs carrying a natural (2′-deoxy)ribose are quite exceptional since, apart from the DNA virus inhibitor brivudin (and related 5-substituted analogs) and the RNA virus inhibitors favipiravir (in activated form), molnupiravir and ribavirin, no approved examples exist [12]. We were therefore intrigued to understand the precise mechanism underlying EDA’s inhibitory effect on CoV replication. Our investigation, reported herein, covered assessment of EDA across multiple cell models and CoVs, including engineered viruses carrying core substitutions in nsp12 or nsp14.

Next, we conducted assays with EDA-triphosphate (EDA-TP) and the SARS-CoV-2 RdRp and ExoN enzymes, to establish the inhibitory mechanism on viral RNA synthesis. The biological findings were corroborated through cryogenic electron microscopy (cryo-EM) of SARS-CoV-2 RdRp in complex with an EDA-terminated RNA. Our findings position EDA as a base-modified yet non-mutagenic NA that interferes with CoV RNA synthesis through a combination of immediate stalling, template-dependent inhibition, and possibly additional downstream effects on viral RNA function. Collectively, this qualifies EDA as a mechanistically distinct NA prototype having potential relevance for clinical development.

## Results

### The 6-ethyl function is key to achieve robust activity across multiple CoV cell models

Human coronavirus 229E (HCoV-229E)-infected human embryonic lung fibroblast (HEL299) cells are a convenient model for antiviral screening, since the experiments are done at biosafety level 2 and the virus induces manifest cytopathic effect (CPE) that can be easily quantified by cell viability testing. Using this model, we evaluated a previously synthesized [26, 27] series of eighteen 6-alkyl and 6-aryl substituted 7-deazapurine riboside analogs (S1 Table). The 6-ethyl analog (= EDA; Fig 1A) stood out as a strong inhibitor of HCoV-229E replication (Fig 1B), with an EC_50_ value of 1.8 µM and 50% cytotoxic concentration (CC_50_) >100 µM, resulting in a selectivity index (SI; ratio of CC_50_ to EC_50_) of >56. The 6-methyl analog (MeDA; Fig 1A) was considerably more active but also much more cytotoxic [EC_50_ = 0.036 µM; CC_50_ = 0.64 µM; SI = 18; S1 Table].

Four other analogs showed weak activity (see S1 Table for compound structures), with EC_50_ values of 26 µM (R = cyclohexyl; compound **15**); 35 µM (R = 4-ethylphenyl; compound **10**); 47 µM (R = isopropyl; compound **3**); and 53 µM (R = propyl; compound **2**). We next evaluated nine analogs of EDA, bearing a short heteroatom-substituted alkyl group (S2 Table). No anti-HCoV-229E activity was seen for three isosteric analogs of EDA bearing a thiomethyl (**17**), methoxy (**18**) or methylamino (**19**) function. The same was true for compounds **21**-**25**, bearing a heteroatom-substituted isopropyl group, whereas the 2-hydroxyethyl (**20**) analog showed an EC_50_ value of 15 µM (S2 Table). This indicates a narrow structure-activity relationship with a key role for the 6-ethyl substituent. This structural element proved crucial to achieve antiviral effect at non-toxic concentrations, since unmodified 7-deazaadenosine (= tubercidin) had an EC_50_ of 0.29 µM and CC_50_ of 2.0 µM, yielding a SI of only 7 (S1 Table). EDA showed comparable micromolar potency as the reference compound GS-441524 (= nucleoside form of RDV; EC_50_: 2.5 µM), while *N*^4^-hydroxycytidine (= nucleoside form of molnupiravir; NHC) was ∼3-fold more potent (S1 Table). The strong inhibition of HCoV-229E replication was confirmed when we analyzed the supernatants for viral load, using RT-qPCR, or for infectious virus titer (Fig 1C). Both parameters were reduced by 10,000-fold at 14 µM EDA and 7 µM GS-441524, while the EC_99_ value (= concentration to reduce the viral load by 100-fold) was 4.4 µM for EDA and 3.7 µM for GS-441524.

Broader evaluation validated EDA as a robust inhibitor across different α- and β-CoVs and cell models. Our virus panel contained the β-CoVs SARS-CoV-2 (Fig 1E-H) and HCoV-OC43, as well as feline infectious peritonitis virus (FIPV) which, alike HCoV-229E, belongs to the α-genus (Fig 1H). We included four human respiratory epithelium cell models, namely 16HBE cells infected with HCoV-229E (Fig 1D); and Calu-3 cells, A549^ACE2/TMPRSS2^ cells and HNECs grown at the air-liquid interface, each infected with SARS-CoV-2 (Fig 1E-H). As summarized in Fig 1H, the average values for EC_50_ (CPE or dsRNA staining) or EC_99_ (viral load) were in the range of 1 to 10 µM for EDA, 0.01 to 0.09 µM for MeDA, 0.6 to 12 µM for GS-441524 and 0.2 to 6 µM for NHC. This broad testing confirmed that EDA has a markedly higher SI value than MeDA, pointing to the 6-ethyl group as a key substituent for achieving robust activity combined with a favorable therapeutic window.

### The CoV inhibition by EDA does not involve virus mutagenesis

To obtain a first insight in the mechanism of EDA, we conducted a one-cycle time-of-addition assay [30] in HCoV-229E-infected HEL299 cells, in which EDA or GS-441524 were added at different time points relative to the time of virus infection. At 16 h p.i., cell extracts were prepared to quantify the viral RNA by RT-qPCR. Whereas the entry inhibitor E64d lost most of its activity when added at 2 h p.i., EDA and GS-441524 showed a very similar profile in maintaining strong activity when added up until 8 h p.i., to then lose activity afterwards (S1 Fig). This was a first indication that EDA acts at the stage of viral RNA synthesis, which aligns with inhibition of the viral polymerase.

Considering that EDA carries an unnatural purine base combined with a natural ribose, we wondered whether it might be incorporated into the viral genome and induce a mutagenic effect on the virus. To address this, HCoV-229E was passaged in HEL299 cells in the presence of 1.3 µM EDA, GS-441524 or NHC (Fig 2A). After the fourth passage, the viruses were submitted to whole genome Nanopore sequencing. To avoid that the error rate of this methodology could skew the analysis, we applied a threshold of 10% to calculate the number of mutations under each drug regimen. The virus produced under NHC carried in total 69 mutations and, as expected [7, 31], the majority of the mutations were C-to-U, U-to-C and G-to-A transitions (Fig 2B). In sharp contrast, only one mutation (G-to-U) was present in the GS-441524 condition and none was detected in the virus passaged under EDA. Accordingly, NHC caused a gradual loss in infectious titer, with virus extinction at passage #5 (Fig 2C). For the conditions receiving EDA, GS-441524 or no compound, the infectious titer remained stable over the five passages. Hence, alike GS-441524, EDA does not induce (lethal) virus mutagenesis.

**Fig 2.**
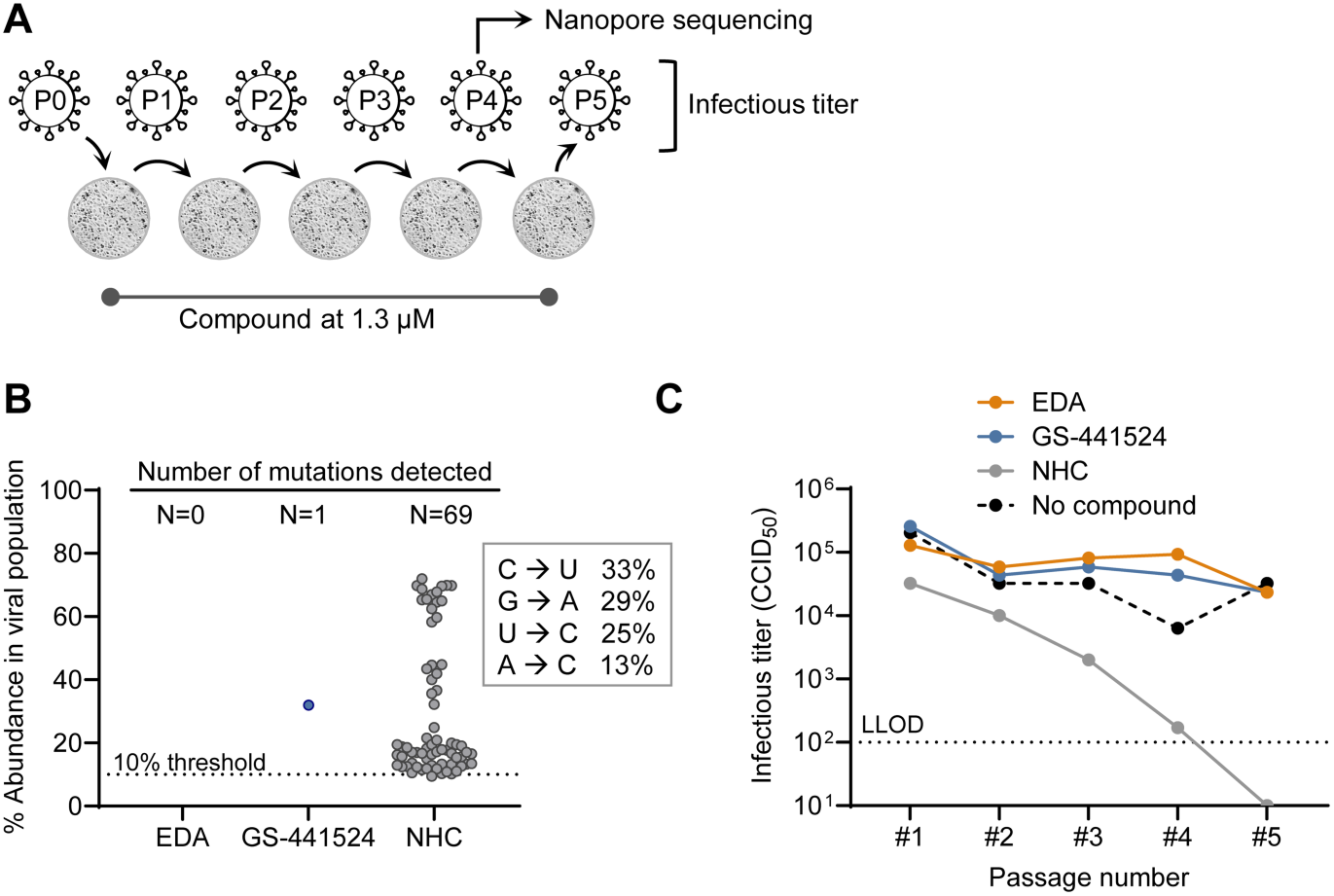
EDA does not act as a virus mutagen. (**A**) HCoV-229E was passaged five times in HEL299 cells, under 1.3 µM of EDA, GS-441524 or NHC. (**B**) Number and abundance of mutations detected by Nanopore sequencing of the #4 viruses, applying a threshold of 10% in the total number of reads. On the right, the proportion of each mutation type, detected in the NHC condition, is specified. (**C**) At each passage, the harvested viruses were analyzed for infectious virus titer. LLOD: lower limit of detection. The data shown are from a single passaging experiment.

### Mutations in the RdRp core of SARS-CoV-2 nsp12 render the virus less sensitive to EDA

To next validate our assumption that EDA targets the viral RdRp, we evaluated the compound against engineered SARS-CoV-2 viruses bearing a selection of mutations in the RdRp core of nsp12. The panel included single mutants E802D^nsp12^ and C799F^nsp12^, and double mutant (S759A+V792I)^nsp12^, associated with mild to prominent resistance to RDV. These mutations emerged after passaging of SARS-CoV-2 under GS-441524 or RDV [32] or during administration to COVID-19 patients of RDV or obeldesivir, the oral prodrug of GS-441524 [33–37]. Mutation S759A is located in the active site motif C, while the other three are clustered in or adjacent to motif D, in the vicinity of the active site [32]. Considering the presumed relevance of motif D for the inhibition by EDA (as apparent from the cryo-EM analysis; see last part of the Results section), we analyzed the ∼17 million SARS-CoV-2 sequences in the GISAID database for variation in this part of nsp12. We found that Ala797 exhibits natural variation, albeit at very low frequency, with substitutions to Glu, Ser, Thr or Val. A797 lies adjacent to K798, which is strictly conserved and crucial for interaction with the γ-phosphate of the incoming NTP [38]. Based on this, we decided to also engineer mutant viruses bearing A797E^nsp12^, A797S^nsp12^, A797T^nsp12^ or A797V^nsp12^.

Upon evaluation in A549^ACE2/TMPRSS2^ cells, the A797T^nsp12^-, C799F^nsp12^- and E802D^nsp12^-mutant viruses exhibited ∼3-fold lower sensitivity to EDA, showing EC_99_ values of 5.9, 6.0 and 4.5 µM, compared to 1.7 µM for WT virus (Fig 3). Other substitutions at residue 797 had lower impact on the activity of EDA; this was most clear for A797E^nsp12^ for which the EC_99_ value was the same as for WT. For GS-441524, the C799F^nsp12^- and E802D^nsp12^-mutants showed about 6- and 3-fold resistance, respectively, in agreement with other reports on GS-441524, RDV or obeldesivir [32–35, 39]. A clear difference between the two antiviral molecules was evident for the double mutant (S759A+V792I)^nsp12^ which, compared to WT virus, showed 14-fold lower sensitivity to GS-441524, while its EC_99_ value for EDA was not significantly different from that of WT virus. For NHC, all nsp12 mutants had a comparable EC_99_ value as WT virus; the A797T^nsp12^ virus exhibited a minor increase (i.e. 1.8-fold) which reached significance. This indicates that the tested mutations may not influence the activity of an NA that does not cause stalling of RNA synthesis [40].

**Fig 3.**
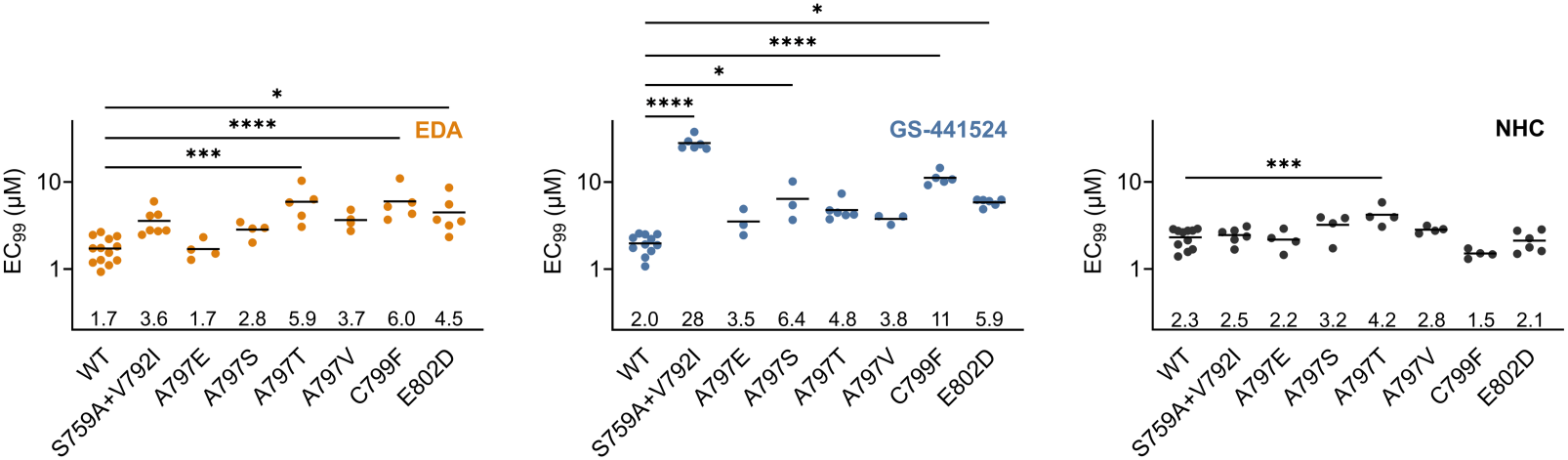
Evaluation against engineered SARS-CoV-2 viruses bearing mutations in nsp12. The S759A+V792I, C799F and E802D substitutions confer moderate resistance to GS-441524, and have been identified after RDV or obeldesivir treatment of SARS-CoV-2-infected cells [32] or patients [33–37]. Four additional substitutions were engineered at A797, a residue adjacent to the conserved K798 residue that is critical for binding of the incoming nucleotide [38]. The Y-axis shows EC_99_ values determined by RT-qPCR for viral load. The dot symbols show individual data points from 3-13 independent experiments. Mean values are shown as bars and are also specified above the X-axis. Mutants were compared with wild type (WT) by ordinary one-way ANOVA with Dunnett’s multiple-comparisons correction.

### The anti-CoV activity of EDA is impeded by the ExoN function of nsp14

Excision by the nsp14-ExoN may reduce the anti-CoV activity of NAs, with compounds such as ribavirin and 5-fluorouracil showing far greater susceptibility to excision than RDV and NHC [7, 23, 24]. A convenient way to study this in cell culture is to use a mutant CoV lacking functional ExoN activity due to substitutions at catalytic residues. So far, the rescue of replication-competent ExoN-deficient virus has been successful for mouse hepatitis virus (MHV) [41] and SARS-CoV [22], but not for MERS-CoV and HCoV-229E [42, 43]. In two recent studies, ExoN-deficient SARS-CoV-2 mutant viruses were successfully rescued, but displayed severely impaired replication fitness [44, 45]. This potentially explains why no viable virus was rescued in an earlier investigation [42] as well as our own attempt to reverse-engineer a SARS-CoV-2 mutant carrying the catalytic D90A and E92A substitutions.

To circumvent this, we engineered two other SARS-CoV-2 mutants bearing a substitution at a non-catalytic nsp14 residue. Substitution P203L^nsp14^ lies within an nsp14-nsp10 interaction domain and was suggested to reduce the SARS-CoV-2 ExoN proofreading activity, based on the higher mutation rate observed in hamsters infected with an engineered P203L^nsp14^ virus [46]. Residue Q145 lies within the ExoN catalytic core [25, 47] and was proposed to interact with the base of the 3′-terminal nucleotide in nascent RNA [25]. Since a low number of the SARS-CoV-2 sequences in the GISAID database bear an Arg at this position, we reasoned that a Q145R^nsp14^ mutant virus might be viable, which indeed proved to be the case. Upon antiviral testing, the activity of EDA, GS-441524 and NHC was not significantly affected by either substitution P203L^nsp14^ or Q145R^nsp14^ (S2 Fig). This suggests that Q145 may not play a major role in mediating excision of EDA, or that substitution with R145 does not disrupt this function. For P203L^nsp14^, our data are consistent with the report that this mutant displayed an increased mutation rate in hamsters, but not in cell culture [46].

Because a robust replication-competent SARS-CoV-2 mutant lacking ExoN activity was not available, we next used MHV as a surrogate model. Mouse L2 cells were infected with either WT MHV or an nsp14-mutant carrying substitutions D89A and E91A, which render the virus ExoN-deficient. As a reference compound expected not to be affected by this deficiency, we used *Urtica dioica* agglutinin (UDA), a lectin inhibitor of viral entry [48]. Indeed, the dose-response curves of UDA (Fig 4) showed complete overlap for mutant and WT. For EDA, the curve of the mutant showed a prominent shift to the left and, as a result, its EC_99_ was 65-fold lower for ExoN-deficient than for WT virus. Hence, the anti-MHV effect of EDA seems severely impeded by ExoN excision. Nevertheless, EDA exhibited lower potency against WT MHV (EC_99_ of 26 µM) compared to HCoV-229E, SARS-CoV-2, OC43 and FIPV, as evident from the EC_50_ and EC_99_ values for these four viruses, described above. Among multiple factors that might explain this difference, we speculate that mouse L2 cells may convert EDA less efficiently into its active 5′-triphosphate metabolite, similar to the interspecies dependency seen with other adenosine analogs [49].

**Fig 4.**
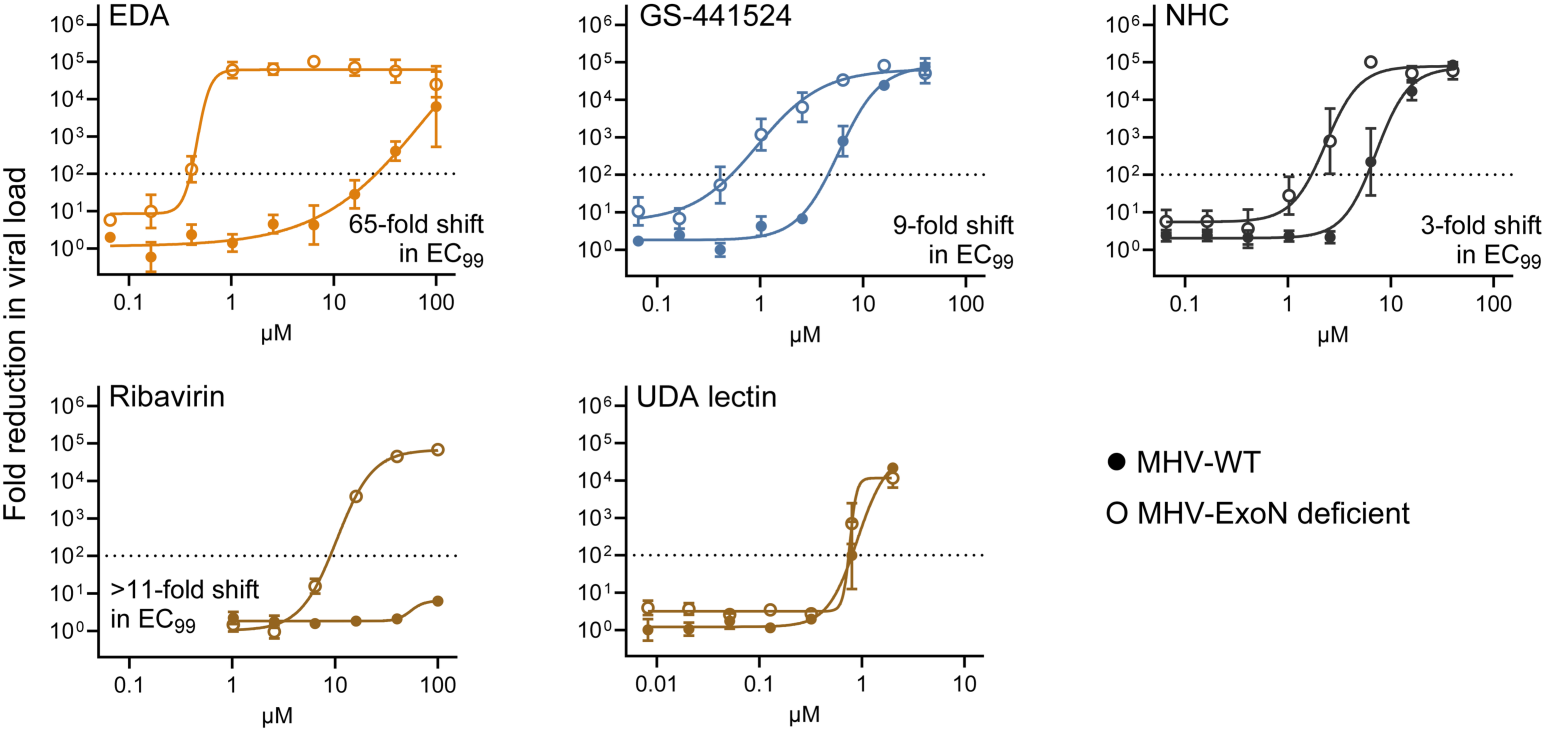
EDA is markedly more active when the virus lacks a functional ExoN proofreading enzyme. Dose-response curves for EDA and four reference compounds, for inhibition of WT or ExoN-deficient MHV in mouse L2 cells. The Y-axis shows the fold reduction in viral load, based on RT-qPCR. Data are the mean ± SEM (N=3). For UDA lectin, an inhibitor of viral entry, the activity is not affected by ExoN deficiency. For the other compounds, the fold reduction in EC_99_ value between ExoN-deficient and WT virus is specified in the graph.

For GS-441524, ExoN-deficient MHV exhibited a 9-fold increase in sensitivity compared with WT virus, whereas NHC showed a 3-fold shift (Fig 4). This aligns with two other studies evaluating RDV and NHC against ExoN-deficient MHV [7, 24]. Likewise, we confirmed that ExoN deficiency renders MHV highly sensitive to ribavirin [23], which showed an EC_99_ of 9 µM for mutant MHV while having no effect on the WT virus at a concentration as high as 100 µM.

In summary, our cell-based virus inhibition studies identify the 6-ethyl moiety of EDA as a key element for robust and selective anti-CoV activity across multiple CoVs and cell models. EDA exerts its antiviral effect at the stage of viral RNA synthesis by targeting the RdRp complex, as supported by its diminished activity against nsp12-mutant forms of SARS-CoV-2 and its increased potency against ExoN-deficient MHV.

### EDA is incorporated by SARS-CoV-2 RdRp but also sensitive to excision by ExoN

To learn how EDA precisely interferes with viral RNA synthesis, we conducted enzymatic assays with EDA-5′-triphosphate (EDA-TP), the SARS-CoV-2 RdRp and ExoN enzymes, and different variations of the natural NTPs and RNA hairpin or primer:template substrates.

We first assessed the concentration dependency for single nucleotide incorporation by the RdRp using a fixed reaction time of 30 s. Both ATP and EDA-TP produced concentration-dependent primer extension, although the EDA-TP curve was shifted towards higher nucleotide concentrations. Accordingly, the nucleotide concentrations yielding half-maximal product formation after 30 s were 0.74 µM for EDA-TP *versus* 0.13 µM for ATP (Fig 5A). In a time-course experiment with a different primer:template system, EDA-TP was incorporated at approximately half the rate of ATP at 0.4 µM nucleotide concentration (S3 Fig), while at 2 µM, the incorporation rates of EDA-TP and ATP were similar. In line with the lack of mutagenic effect in CoV-infected cells (see above), EDA-TP did not incorporate opposite to C, G or A, even after 60 min reaction time with 20 µM EDA-TP (S4 Fig). When the EDA-TP concentration was increased to 200 µM, a barely detectable band was observed for its incorporation as a mimic of GTP or UTP (S4 Fig).

**Fig 5.**
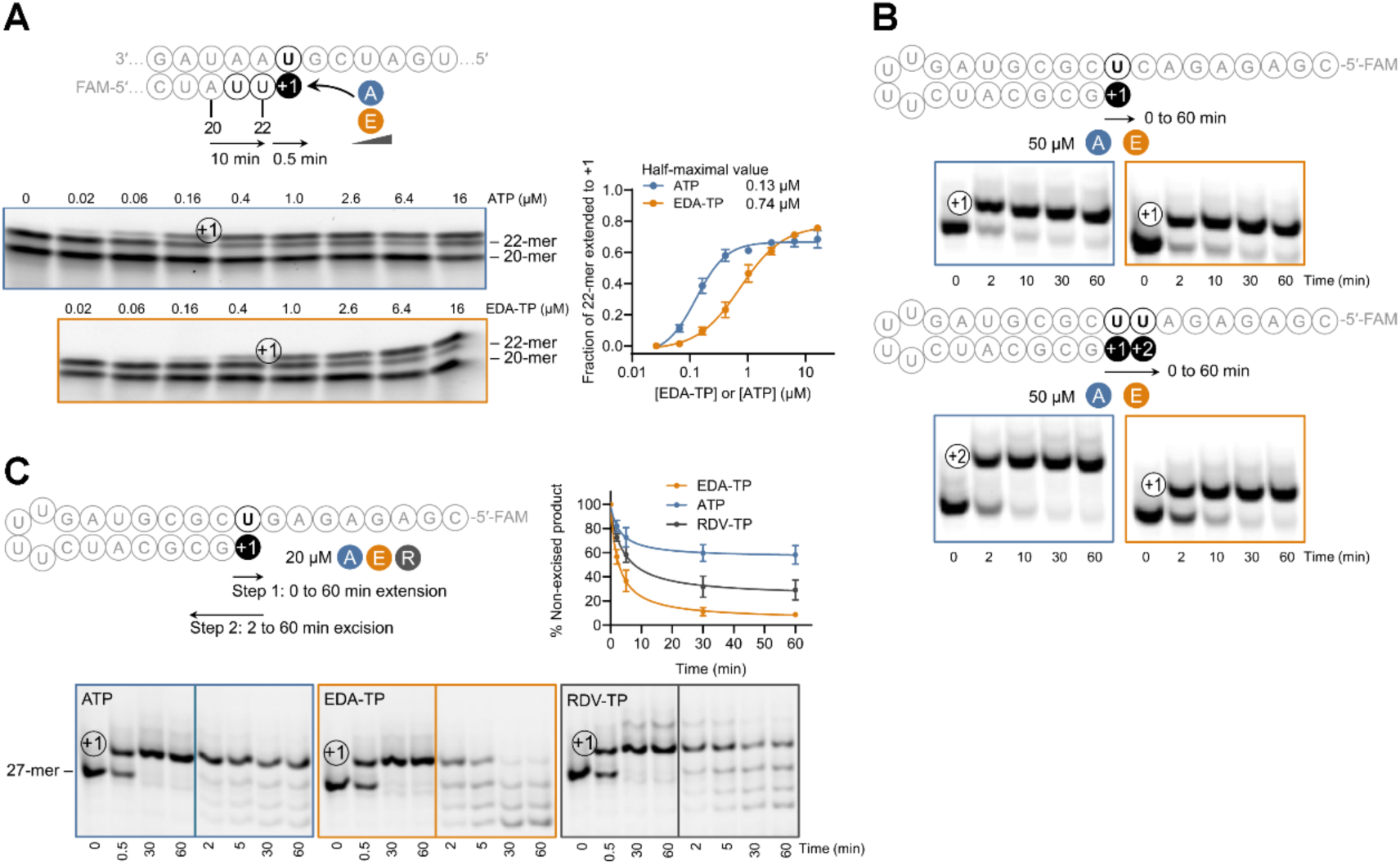
EDA incorporation by SARS-CoV-2 RdRp and susceptibility of EDA-terminated RNA to ExoN-mediated cleavage. (**A**) Single nucleotide incorporation by SARS-CoV-2 RdRp, using different concentrations of EDA-TP or ATP, and 30 s reaction time. The plot shows the fraction of the 22-mer that was extended with EDA-TP or ATP (mean ± SEM; N=3). GraphPad Prism was used for nonlinear regression curve fitting and for calculation of the half-maximal values, i.e. nucleotide concentrations yielding half-maximal product formation at the 30 s endpoint. (**B**) Incorporation of EDA-TP or ATP, at a concentration of 50 µM and monitored over time. Top: hairpin template with one U. Bottom: against a hairpin template with two consecutive Us, the incorporation of EDA-TP stops at +1 and disables addition of EDA-TP at +2. (**C**) Time-course for ExoN cleavage of RNA products terminating in EDA, A or RDV following sequential RdRp extension and ExoN excision. After 60 min polymerization in the presence of EDA-TP, ATP or RDV-TP, the RdRp was inactivated and the ExoN was added. Graph: mean ± SEM from N=3.

The efficient incorporation of EDA-TP as a substitute for ATP was also seen using a hairpin substrate. At 50 µM of EDA-TP or ATP, extension to the +1 product was complete within 2 min reaction time (Fig 5B). Intriguingly, the use of a hairpin with two consecutive U residues showed that the incorporation of EDA-TP at position +1 does not allow any extension with a second EDA-TP at +2. Even after 60 min incubation with 50 µM EDA-TP, no +2 product was detected, which sharply contrasted with the incorporation of two consecutive ATPs being complete after 2 min (Fig 5B).

Next, we conducted a sequential extension-excision assay in which we first allowed 100% extension of the hairpin RNA with a single EDA-TP, ATP or remdesivir-5′-triphosphate (RDV-TP), after which the resulting +1 products were exposed to SARS-CoV-2 ExoN. Under these assay conditions, 58% of the ATP-extended product was still intact after 60 min incubation with ExoN (Fig 5C). The EDA-TP-extended product declined to 57% after only 2 min, and to 9% after 60 min, whereas the RDV-TP-extended product showed an intermediate excision profile (Fig 5C). Thus, under the conditions tested, EDA-terminated RNA was more susceptible to ExoN-mediated cleavage than the ATP- or RDV-terminated products.

### EDA causes immediate stalling which is transient and also occurs against templating EDA

As mentioned above, the RdRp appeared incapable of extending an EDA residue with a second EDA-TP. Chain extension with a natural NTP was still possible but also compromised. Under short reaction conditions (30 s), CTP incorporation was less efficient after EDA than after A (Fig 6A). While the latter reaction achieved already its plateau of 94% at 0.8 µM CTP, the incorporation of CTP after EDA did not surpass 68%, despite the use of up to 100 µM CTP.

**Fig 6.**
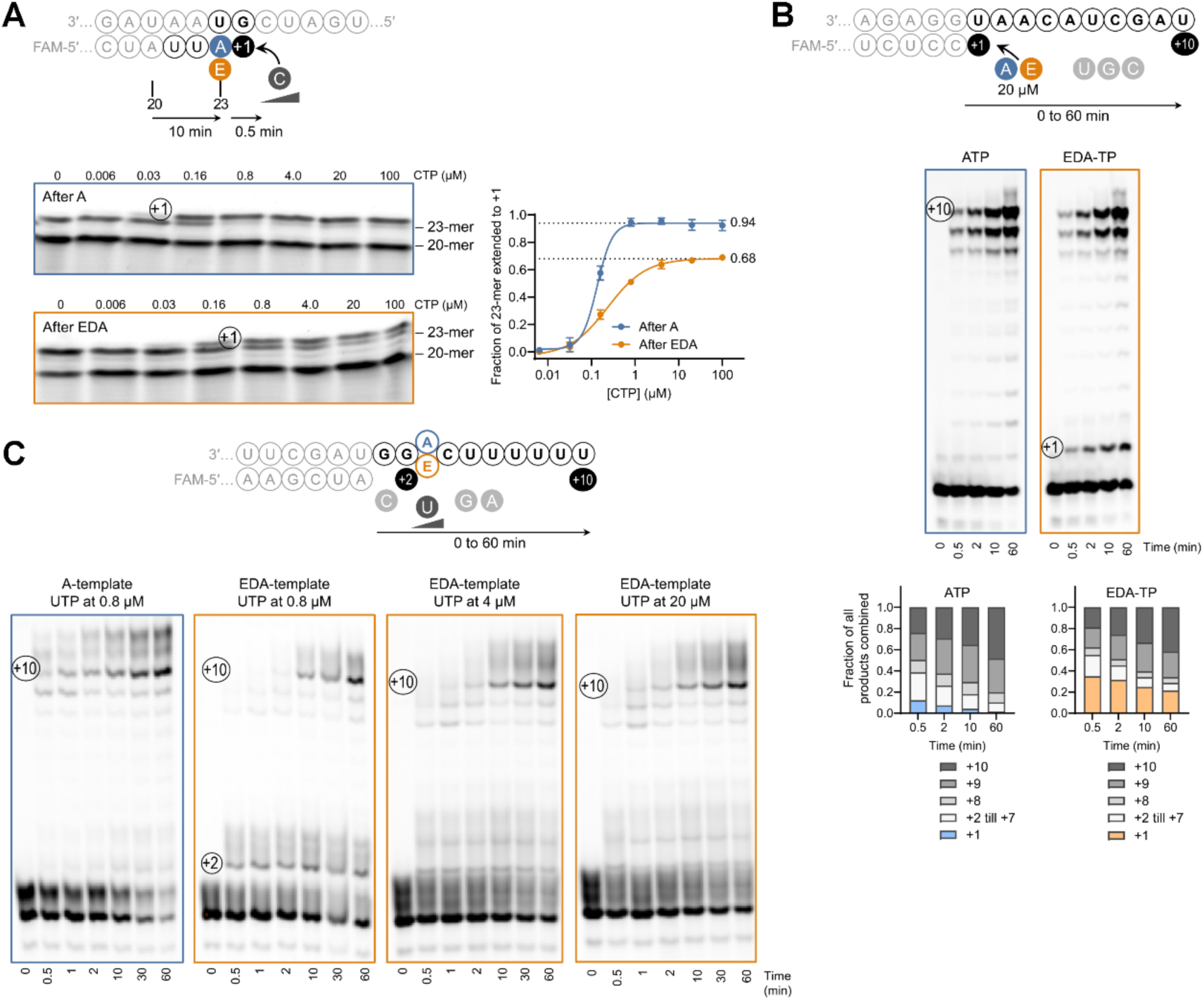
Post-incorporation and template-dependent stalling by EDA. (**A**) The incorporated EDA impairs extension with the next NTP. Graph: fraction of the 23-mer, ending in A or EDA, that was extended with CTP. Data points are the mean ± SEM (N=3), with curves fitted by nonlinear regression. The fitted plateau values are specified next to the dotted lines. (**B**) The stalling effect is transient and still permits further chain extension. Graph: relative band intensities (mean; N=2) of the +1 intermediate with incorporated ATP (blue) or EDA-TP (orange), and of the longer products (different shades of grey). (**C**) Stalling induced by a templating EDA, as evident from transient accumulation of the +2 intermediate that precedes +3 incorporation of UTP opposite to EDA. The image shows the 0.8, 4 and 20 µM UTP conditions assayed against the EDA-template, compared to the 0.8 µM UTP condition against the A-template. See S5 Fig for additional data with the A-template and 100 µM UTP, and for a plot of the band intensities.

The impaired extension beyond EDA proved not permanent and quite readily overcome. This was evident when we monitored the incorporation of ten nucleotides, starting with EDA-TP or ATP at +1 (Fig 6B). Throughout the 0.5-60 min time course, the +1 product with 3′-terminal EDA showed only a slow decrease, representing 35% of the total of all products at 0.5 min and remaining at 22% after 60 min. For the +1 product with 3′-terminal A, the corresponding values were 5% and 1%. On the other hand, the incorporated EDA did not prevent extension to the final +10 product, which made up 42% of the total at 60 min, compared to 48% in the condition with incorporated A.

The above findings suggest that virus replication in the presence of EDA may lead to (sub)genomic viral RNA with incorporated EDA, resulting in a potential effect on next-round RNA synthesis. To address this, we conducted an assay with an in-house synthesized template containing a single EDA residue (see partial sequence in Fig 6C and full sequence in S3 Table). Whereas the A-containing template allowed for uninterrupted extension to the longer (= +8, +9 and +10) products, the EDA-containing template induced accumulation of a short +2 product (= primer extended with two C residues; Fig 6C), due to inefficient incorporation of UTP at the +3 site opposite to EDA. This stalling effect of templating EDA was particularly pronounced at short reaction times with 0.8 µM UTP, and partially overcome at later reaction times or UTP concentrations of 4 µM, 20 µM (Fig 6C) or 100 µM (S5 Fig, panel A). Even then, in the EDA-template condition incubated with 100 µM UTP, the fraction of fully extended +10 product was lower throughout the 0.5-60 min time course, compared to that obtained with the A-template (see graph in S5 Fig, panel B).

To conclude, as an efficient alternative substrate for ATP, EDA-TP is readily incorporated by SARS-CoV-2 RdRp but then causes ‘immediate’ stalling of further chain elongation. The stalling effect is not permanent, allows considerable readthrough, and also occurs during polymerization against a template with embedded EDA. Still, compared to ATP and RDV-TP, EDA exhibits higher susceptibility to ExoN-mediated excision, which may compromise its ability to inhibit viral RNA synthesis.

### Cryo-EM analysis reveals that EDA incorporation impairs stable binding of the next NTP

To define the structural basis for the stalling effect of EDA, we determined the cryo-EM structure of the SARS-CoV-2 replication-transcription complex (RTC) bound to a template:primer RNA duplex with 3′-terminal EDA in the primer (Fig 7A,C). By using two different templates, we generated two structures with a variation at the next templating base (i.e. +1U in RTC1:EDA and +1G in RTC2:EDA). The complex contains nsp12, nsp7, two nsp8 and nsp13, and one nsp9, assembled on a 40-bp template:primer RNA duplex. The duplex is accommodated within the canonical cleft formed by the fingers, palm and thumb domains of nsp12, and is further stabilized by nsp8. Consistent with previous RTC structures, extensive contacts are formed between nsp12 and the RNA scaffold, with 43 nsp12 residues directly engaged in RNA binding. Clear density is observed for EDA, which is covalently incorporated at the 3′ end of the primer strand and occupies the −1 position of the product RNA, indicating that the analog has been successfully added to the nascent strand and that translocation has occurred (Fig 7A-D). However, regardless of whether the +1 position of the template strand is U or G, the NTP-binding pocket remains empty, with no incoming EDA-TP (Fig 7B) nor CTP (Fig 7D) being bound.

**Fig 7.**
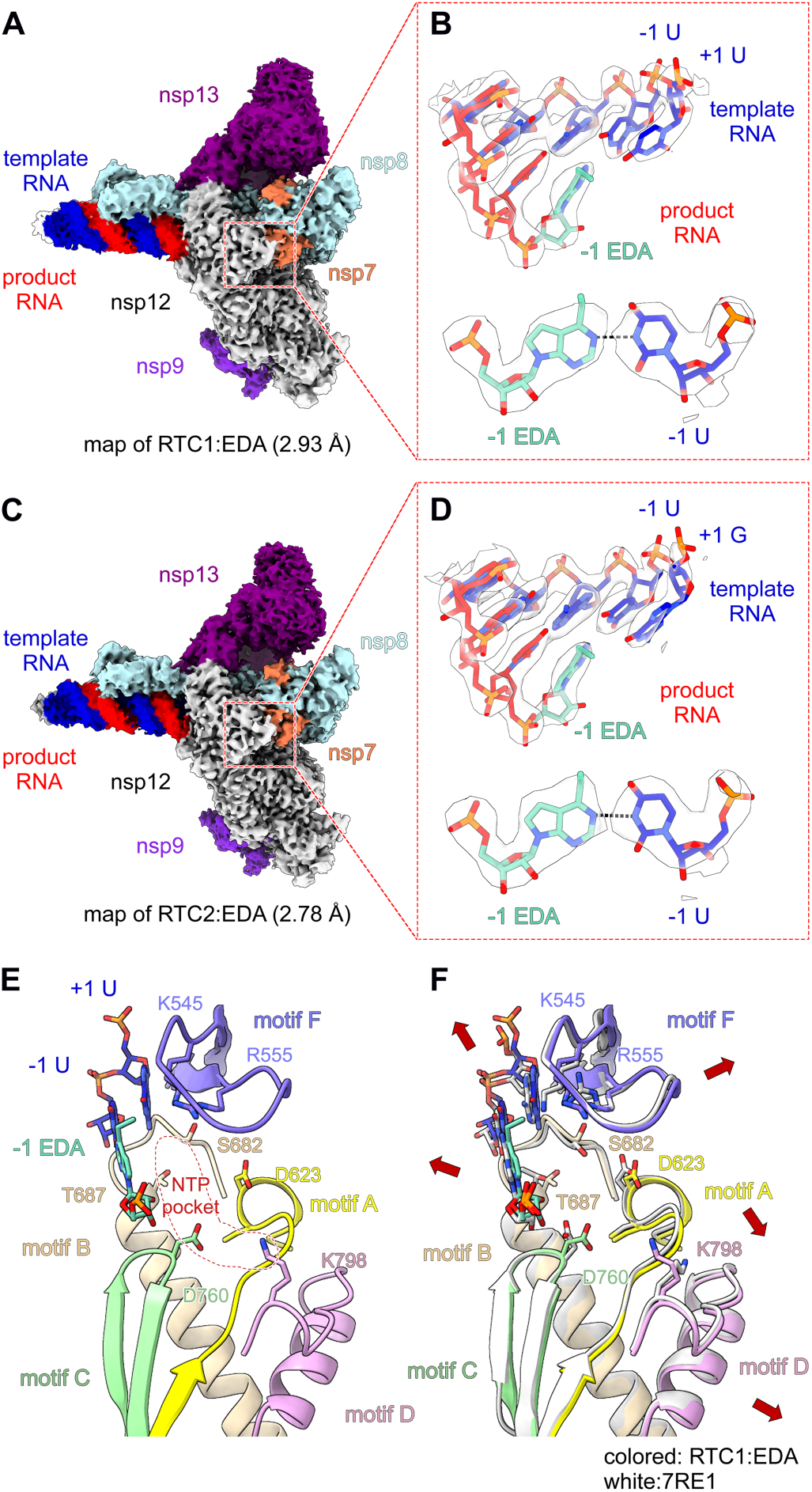
Cryo-EM structure of the RTC:EDA complex. (**A**, **B**) Cryo-EM density of the 2.93 Å resolution RTC1:EDA structure showing, in panel B, the positioning of the incorporated EDA and the nearby bases of the template RNA1 and product RNA strands. (**C**, **D**) Cryo-EM density of the 2.78 Å resolution RTC2:EDA structure showing, in panel D, the positioning of the incorporated EDA and the nearby bases of the template RNA2 and product RNA strands. (**E**) Detailed view of the active site of the RTC1:EDA complex, depicting the spatial organization of conserved RdRp motifs A-D and F (colored yellow, wheat, pale green, pink and light blue, respectively) and of key residues, with the vacant NTP-binding site delineated by a dashed red contour. (**F**) Superposition of the RTC1:EDA active site with that of the apo structure [white, PDB: 7RE1 [50]], revealing conformational changes in specific motifs and residues (indicated by red arrows). Concerted movements in motifs A, D and F result in opening of the active site.

In both the RTC1:EDA and RTC2:EDA complexes, the incorporated EDA is well positioned within the core of the polymerase active site (Fig 7B,D). As an adenosine analog lacking the exocyclic amino group, EDA maintains stacking interactions with the upstream primer nucleotide but forms only a single hydrogen bond with the opposing templating uridine. Its ribose moiety is coordinated by active site residues S759, D761, and C813, while S814 contacts the phosphate backbone, together stabilizing the incorporated analog within the catalytic center. Thus, EDA can be accommodated after incorporation without gross distortion of the RNA duplex.

Given the overall similarity between the RTC1:EDA and RTC2:EDA complexes, we next selected RTC1:EDA for comparison with previously reported structures [50] representing post-translocated and incoming NTP-bound states of RTC, to assess how EDA affects the next round of nucleotide addition. Although EDA is accommodated at the −1 position, its incorporation induces subtle but functionally important rearrangements in the active site, particularly within motifs A-D and F, resulting in a slightly more open NTP-binding pocket (Fig 7E). Notably, residues K545, R555, D623, S682, T687, D760 and K798, which are critical for coordinating and positioning the incoming NTP [38, 51, 52], adopt outward-shifted side-chain conformations in the EDA-containing complex (Fig 7E,F). These changes would be expected to weaken stabilization of the next nucleotide and reduce the efficiency of productive NTP or EDA-TP binding. Specifically, residues K545 and R555, located in motif F, play a crucial role in positioning the NTP substrate for catalysis, by interacting with both the nucleobase and the triphosphate groups. K545 selectively engages with the incoming NTP, while R555 either forms a pi-pi stacking interaction with the incoming ATP substrate base or establishes an H-bond with the β-phosphate oxygens of incoming GTP, CTP, or UTP. S682 and T687 in motif B, along with D760 in motif C, form a network of polar contacts with the ribose of the incoming NTP. Finally, K798 in motif D interacts with the γ-phosphate of the NTP. When these residues adopt outward-shifted conformations, the stability of the bound NTP in the active site is likely diminished, which could impair subsequent nucleotide addition.

These structural observations provide a mechanistic explanation for our biochemical data. Whereas the incorporation of a second EDA-TP is abolished and the addition of a natural NTP is clearly impaired, no stable density corresponding to the next incoming nucleotide is observed in both the RTC1:EDA and RTC2:EDA structure. Though not acting as a bona fide chain terminator, EDA appears to perturb the geometry of the NTP-binding pocket following its incorporation, thereby impairing the accommodation of the next nucleotide or nucleotide analog and stalling further chain extension.

The proposed mechanism may also explain why SARS-CoV-2 is less sensitive to EDA when bearing nsp12 substitutions A797T, C799F or E802D. All three are located in motif D, a region not directly responsible for NTP coordination but critical for shaping the active site and supporting nucleotide addition. Substitutions at these positions may alter the conformational flexibility of motif D, thereby reducing the recognition or incorporation of EDA-TP, or compensating for the EDA-induced relaxation of the NTP-binding pocket after incorporation of the compound. In either case, such mutations could reduce the extent to which EDA disrupts subsequent nucleotide addition. In contrast, substitution S759A (in combination with V792I) caused prominent resistance to RDV while not changing the activity of EDA. S759 is part of the catalytic SDD motif and its substitution with Ala was shown to reduce the discrimination of RDV-TP relative to ATP [32] and to decrease the catalytic rate by altering the conformation of the ribose at the 3′-end of the primer [53].

## Discussion

Although NAs are among the earliest known antiviral agents, their structural and mechanistic versatility continues to drive the development of new antiviral therapeutics which find their way to the clinic [11–13, 54]. In this study, we elucidated the mechanistic basis for the anti-CoV activity of EDA, a recently discovered [27, 28] analog with modified base yet natural ribose. Our findings support a model in which this 6-ethyl-modified adenosine analog is smoothly incorporated by the CoV RdRp, to then disturb the synthesis or functionality of viral RNA, leading to inhibition of viral replication.

The 6-ethyl group of EDA emerges as a key element for achieving robust and selective anti-CoV activity. This is evident from our antiviral data on a series of analogs differing in size or heteroatom composition of the substituent at position 6. EDA is markedly more selective than its tubercidin congener (= 7-deazaadenosine) or MeDA, the analog bearing a 6-methyl group. The latter was previously reported to have antiviral activity against polio and dengue viruses [55]; hepatitis C replicon [56]; and influenza virus [57], but in each of these as well as our study, MeDA had pronounced cytotoxicity. Although the antiviral potency of EDA may seem rather modest [i.e. EC_50_ values in the micromolar range], this might be readily overcome with suitable protides generating higher intracellular concentrations of the phosphorylated metabolites [58].

Mechanistically, EDA shares certain features with the clinically approved CoV inhibitor RDV; however, it also exhibits clear mechanistic differentiation. Both compounds do not function as bona fide chain terminators of SARS-CoV-2 RdRp, since they both permit considerable readthrough beyond the point at which their stalling effect occurs. Considering that the intracellular NTP concentrations are in the high micromolar to low millimolar range [59], we anticipate that quite some readthrough may occur in CoV-infected cells exposed to EDA. If EDA becomes embedded in viral (sub)genomic RNA or mRNA, it may interfere with downstream RNA synthesis or translation, or some regulatory function of structured viral RNA [29]. Notably, our virus passaging experiment shows that incorporation of the compound does not induce mutations in viral RNA. From a safety perspective, the lack of overt RNA mutagenicity represents a favorable attribute. Enzymatic findings corroborate that EDA-TP is not recognized as a GTP, CTP or UTP mimic, confirming the absence of ambiguous base-pairing. The cryo-EM analysis validates that EDA is efficiently incorporated opposite to U, with favorable pi-pi stacking interactions likely offsetting the formation of a single H-bond rather than two.

Complementary to the above mechanism, EDA impairs viral RNA synthesis by a mechanism referred to as ‘immediate’ stalling, since it occurs right after EDA has been incorporated. At this stalled state, no extension with a second EDA-TP is allowed. The relevance of this observation is not entirely clear, as incorporation of two adjacent EDA residues may be rare given the abundant intracellular levels of ATP. Instead, and consistent with our cryo-EM data, the more probable scenario is that incorporation of EDA impairs extension with the next natural NTP. Concomitantly, this stalled state may trigger excision of EDA by the ExoN domain of the RTC [60], highlighting an important difference between EDA and RDV. Since the stalling by RDV occurs in a ‘delayed’ manner after the addition of three more NTPs [14–16], its removal may require substantial backtracking [18] to allow excision of several nucleotides, from the 3′-end up to the RDV incorporation site. Since the stalling by EDA happens ‘immediately’ at the site where it is incorporated, the excision process is plausibly more efficient. The different excision rates of EDA and RDV are evident from our combined RdRp-ExoN enzymatic assay, and are also supported by our antiviral studies with MHV, in which loss of ExoN function increased the sensitivity to EDA by 65-fold, compared with a 9-fold increase for GS-441524. Still, EDA and GS-441524 have similar antiviral potency in cells infected with human CoVs, which aligns with both NAs serving as efficient substrates for the viral RdRp, but seems at odds with the analysis that EDA undergoes much more excision. Although our study was not designed to measure precise enzyme kinetics, it seems to indicate that the efficient use of EDA-TP as a substitute for ATP, combined with the fast nucleotide incorporation rate of coronaviral RdRps [5], may override the excision of EDA. This structurally unique NA could be a valuable tool to explore the dynamics of CoV RNA transcription [61, 62], and kinetic interplay between the RdRp and ExoN.

The immediate stalling by EDA is well corroborated by our cryo-EM data, which support a model in which EDA acts through a post-incorporation mechanism. Although EDA is incorporated at the 3′-end of the primer strand and well-accommodated at the −1 position, its 6-ethyl modified base induces subtle rearrangements in RdRp motifs A-D and F, leading to outward-shifted conformations for several residues involved in binding of the incoming NTP and, hence, a slightly more open NTP-binding pocket. These changes are expected to reduce the stability of the incoming NTP within the active site and impair nucleotide addition. This model is consistent with our biochemical observations showing that incorporation of a second EDA-TP is abolished, whereas the next natural NTP can still be incorporated at reduced efficiency. The absence of an incoming EDA-TP in the RTC1:EDA complex and of a natural NTP in the RTC2:EDA complex suggests that, in both cases, the nucleotide binding may be transient or too weak to allow capturing of a stably bound state. Our structural model also provides an explanation for the partial EDA resistance of mutant viruses bearing substitutions in nsp12 motif D. While the present study focused mainly on nsp12 mutations selected under RDV, a follow-up study is underway to isolate and characterize mutant viruses selected under EDA. These mutants will further facilitate experimental validation of the RTC:EDA cryo-EM structures and the precise contribution of the different motifs surrounding the active site.

Taken together, our findings position EDA as a robust non-mutagenic inhibitor of CoV replication, acting via a combined mechanism of immediate stalling, template-dependent inhibition and possibly additional downstream effects on viral RNA function. Beyond its relevance for clinical development, EDA exemplifies the ever-expanding potential of NAs as a source of antiviral agents with diverse and distinctive mechanisms of action.

## Materials and methods

### Compounds

The following compounds were purchased from a commercial supplier: GS-441524 and tubercidin (Carbosynth); *N*^4^-hydroxycytidine (NHC; MedChemExpress); remdesivir-5′-triphosphate (SigutLabs); ribavirin (Sigma-Aldrich); E64d (MedChemExpress); and *Urtica dioica* agglutinin (UDA; EY Laboratories). For EDA, MeDA and related 6-alkyl or 6-aryl substituted analogs, the chemical synthesis has been reported elsewhere [26, 27].

### Cells, media and viruses

For human embryonic lung fibroblast cells [HEL299; American Type Culture Collection (ATCC) No. CCL-137], the growth medium contained: Dulbecco’s modified Eagle medium (DMEM); 10% fetal calf serum (FCS); 10 mM HEPES; 1 mM sodium pyruvate; 6 mM L-glutamine; 0.1 mM non-essential amino acids (NEAA); and 25 mM glucose. The infection medium was basically the same, except that Pen-Strep (= 100 U/ml penicillin and 0.1 mg/ml streptomycin) was added and the FCS content was reduced to 2%. Similar growth (10% FCS) and infection media (2% FCS) were used for the Crandell-Rees feline kidney (CRFK) cells (kind gift from H. Nauwynck, UGent); human colon adenocarcinoma HRT-18G cells (ATCC No. CRL-3609); human bronchial epithelial 16HBE cells (kind gift from P. Hoet, KU Leuven); baby hamster kidney BHK-21J cells (donated by P. J. Bredenbeek, Leiden University Medical Center); and mouse L2 fibroblast cells (gift from P. Maes, KU Leuven). Human lung adenocarcinoma Calu-3 cells (ATCC No. HTB-55) were grown in Minimum Essential Medium (MEM) with 10% FCS; 10 mM HEPES; 2 mM L-glutamine; and 0.1 mM NEAA. Finally, human lung carcinoma A549 cells expressing ACE2 and TMPRSS2 (A549^ACE2/TMPRSS2^) were purchased from InvivoGen (No. a549d-cov2r) and cultured in DMEM supplemented with 10% FCS; 6 mM L-glutamine; 25 mM glucose; 100 µg/ml hygromycin and 0.5 µg/ml puromycin. For Calu-3 and A549^ACE2/TMPRSS2^ cells, the infection medium consisted of DMEM with 0.2% FCS; 0.3% bovine serum albumin (BSA); 10 mM HEPES; 1 mM sodium pyruvate; 4 mM L-glutamine; 0.1 mM NEAA; 25 mM glucose; and Pen-Strep. Primary human nasal epithelial cells (HNECs), cultured at the air-liquid interface (ALI), were purchased from Epithelix (pool of 14 donors; EP02MP, MucilAir) and maintained in MucilAir medium (Epithelix, EP04MM). All cells were grown in a humidified 5% CO_2_ incubator at 37 °C.

We used the following viruses: HCoV-229E (ATCC No. VR-740); SARS-CoV-2 early European strain 20A.EU2; reverse-engineered SARS-CoV-2 Omicron BF.7 [wild-type (WT) and nsp12-or nsp14-mutants; see below]; HCoV-OC43 (donated by E. Keyaerts, KU Leuven); feline infectious peritonitis virus (FIPV; donated by H. Nauwynck, UGent); and MHV-A59 virus [i.e. WT or reverse-engineered with mutations D89A and E91A in nsp14; kind donation from V. Thiel (University of Bern)]. Virus stocks were prepared in the same cell line as used for the antiviral assays, then titrated by end-point dilution to define the 50% cell culture infective dose (CCID_50_) via the method of Reed and Muench.

### Reverse-engineering of SARS-CoV-2

Starting from seven plasmids which were generously donated by P.-Y. Shi (World Reference Center for Emerging Viruses and Arboviruses, Texas) [63], we first replaced the sequences corresponding to an early strain of SARS-CoV-2 with the homologous sequences of the Omicron subvariant BF.7. The technical details can be found elsewhere [30]. Depending on the mutation, nsp12 or nsp14 variants were generated either by site-directed mutagenesis of the corresponding plasmids with overlapping primers and Platinum SuperFi II DNA Polymerase (Invitrogen); or by replacing the relevant sequence with a synthetic DNA fragment (Thermo Fisher Scientific) carrying the desired mutation, which was assembled into the PCR-amplified vector backbone (Platinum SuperFi II DNA Polymerase) using the NEBuilder HiFi DNA Assembly Cloning Kit (New England Biolabs). The protocol to generate WT or mutant SARS-CoV-2 was adapted from Xie et al. [63]. In brief, the Omicron BF.7-encoding plasmids were transformed into *E. coli* cells, purified and validated by Sanger sequencing (Macrogen). SARS-CoV-2 DNA fragments were recovered through restriction enzyme digestion of the seven plasmids and subsequent ligation yielded full-length viral DNA. N-gene cDNA was PCR-amplified from the plasmid containing the last part of the viral DNA. The full-length DNA and N-gene DNA were *in vitro* transcribed (mMESSAGE mMACHINE T7 transcription kit; Invitrogen), and the resulting RNA samples were purified by phenol:chloroform extraction and isopropanol precipitation. The genomic RNA and N-coding RNA (20 μg each) were electroporated into eight million BHK-21J cells resuspended in PBS, using the Gene Pulser Xcell Electroporation System (Bio-Rad). Next, the electroporated cells were overlaid on A549^ACE2/TMPRSS2^ cells in a 75-cm² flask. Three days later, the supernatants were harvested and the viruses were expanded in A549^ACE2/TMPRSS2^ cells. The presence of the desired mutations was checked by sequencing with reported primers [63], after which the viruses were titrated and used in antiviral assays.

### Antiviral and cytotoxicity assays

On day 0, serial dilutions of the compounds in medium were added to confluent cell cultures in 96-well plates, immediately followed by the virus at 100 (HCoV-229E, SARS-CoV-2 and HCoV-OC43), 200 (FIPV) or 50 CCID_50_ (MHV-A59) per well. A mock-infected plate was prepared in parallel, to assess compound cytotoxicity. The plates were incubated at 35 °C for HCoV-229E and SARS-CoV-2 in HNECs; and at 37 °C for the other cells and viruses. In the cytopathic effect (CPE) reduction assays, we incubated the cells until day 3 (FIPV in CRFK cells) or day 5 (HCoV-229E in HEL299 cells and SARS-CoV-2 in A549^ACE2/TMPRSS2^ cells) post-infection (p.i.). The compounds’ cytotoxicity and protective effect against viral CPE were monitored by cell viability assay. After adding the CellTiter 96 AQ_ueous_ MTS Reagent (Promega) and incubating for 6 h (A549^ACE2/TMPRSS2^ and CRFK) or 24 h (HEL299 cells) at 37 °C, the OD_490_ values were determined in a microplate reader. The formulas to calculate the compounds’ EC_50_ (half-maximal effective concentration) and CC_50_ (50% cytotoxic concentration) values, are specified elsewhere [64].

For SARS-CoV-2-infected Calu-3 cells, virus replication was quantified at day 3 p.i. by immunostaining for dsRNA. After 15 min fixation with 4% paraformaldehyde; 10 min permeabilization with 0.2% Triton X-100; and 1 h blocking with 1% BSA, the cells were stained with mouse monoclonal anti-dsRNA antibody (J2, SCICONS English & Scientific Consulting Kft) and goat anti-mouse AlexaFluor488 secondary antibody (A21131, Invitrogen), diluted 1:1000 and 1:500, respectively, in PBS with 1% BSA. Nuclei were counterstained with Hoechst (Invitrogen). The percentage of green (infected) cells and the total number of nuclei, as a measure of cytotoxicity, were quantified in a CellInsight CX5 High Content Screening platform (Thermo Fisher Scientific). These readouts were used to calculate the EC_50_ and CC_50_ values from the semi-log dose-response curves (GraphPad Prism).

To monitor virus replication via viral load or titration assay, the cells were infected as above, except that the virus was pre-incubated with the cells for 2 h, to then remove the excess virus and add the compound dilutions. At day 2 (MHV-A59), day 3 (HCoV-229E in 16HBE cells, HCoV-OC43 and SARS-CoV-2 in Calu-3 cells) or day 5 p.i. (HCoV-229E in HEL299 cells and SARS-CoV-2 in A549^ACE2/TMPRSS2^ cells), the supernatants were harvested and frozen at −80 °C. For assays in ALI cultures of primary HNECs, the apical surface was washed on the day of infection to remove mucus. Compounds were added to the basolateral compartment, after which the virus inoculum was applied to the apical side. At 2 h p.i., excess virus was removed by washing. At 24 h p.i., the basolateral medium containing compound was refreshed. Apical washes were collected at 2, 24, 48 and 72 h p.i. and stored at −80 °C.

To determine the number of viral RNA genomes (= viral load), the samples were mixed with Cells-to-cDNA II lysis buffer (Invitrogen) and incubated for 10 min at 75 °C. The one-step RT-qPCR (iTaq Universal Probes One-Step kit, BioRad) employed N-gene specific primers and probes (S4 Table) as previously described [65] and an N-gene plasmid standard customized to the virus. The RT-qPCR program consisted of: 10 min at 50 °C; 2 min at 95 °C; and 40 cycles of 15 s at 95 °C and 60 s at 60 °C. To quantify the titer of infectious HCoV-229E in the supernatant samples, serial dilutions were incubated with HEL299 cells, and the CPE was scored at day 5 p.i. to then calculate the CCID_50_ titer. Antiviral activity was expressed as the EC_99_ value, i.e. compound concentration producing 2-log_10_ reduction in the number of viral RNA genomes or in the CCID_50_ titer.

### Viral fitness and mutational burden after prolonged compound exposure

HEL299 cells were infected with 100 CCID_50_ of HCoV-229E and 2 h later, excess virus was removed and the cells were washed twice before adding the compounds (i.e. EDA, GS-441524 or NHC) at 1.3 µM. At day 4 p.i., the CPE was scored, and the supernatants were frozen at −80 °C. After virus titration, fresh HEL299 cells were exposed to the passaged viruses (100 CCID_50_) and compounds, and the procedure was repeated for a total of five passages. To determine the mutational burden in the #4 passages, we used a similar workflow as described in full detail elsewhere [66, 67] to generate complementary DNA (cDNA) libraries and conduct Nanopore sequencing. Briefly, the supernatants were submitted to RNA extraction using the QIAamp Viral RNA Mini kit (QIAGEN) and the RNA was reverse-transcribed with sol-PrimerA, followed by second-strand synthesis with Sequenase reagents (Thermo Fisher Scientific) and amplification with Sol-PrimerB. For the Nanopore sequencing, sequencing libraries were prepared with the SQK-NBD114.24 kit (Oxford Nanopore Technologies) and 100 ng cDNA of each sample as input. About 80 fmol library was loaded onto an R10.4.1 flow cell. Sequencing was performed on a GridION (Software release 24.02.16), employing super-accurate basecalling and a quality threshold of q>8. CLC Genomics Workbench v22.0.1 was used to trim off primer sequences. Trimmed reads were mapped to the HCoV-229E reference genome (Genbank ID: NC_002645) using minimap2, after which the per-base coverage and relative identity of each position was determined using bam-readcount. Positions in which the fraction of the reference allele was <0.9 were considered mutated sites. Variants shared between all samples, including the original virus stock, were removed from further analyses.

### Enzymatic assays with the RdRp and ExoN proteins of SARS-CoV-2

The enzymatic assays were conducted in two laboratories, where slightly different procedures applied to produce the RdRp complex and evaluate the incorporation of EDA-TP or NTPs towards a primer:template duplex or hairpin RNA, for which the full sequences are provided in Table S3.

### Primer extension and nucleotide excision in function of time

The SARS-CoV-2 (Wuhan-Hu-1 isolate; NCBI sequence YP_009724389.1) nsp7, nsp8, nsp10, nsp12 and nsp14 proteins, in 6His-or 8His-tagged and TEV-cleavable form, were produced in *E. coli*. The proteins were purified by affinity chromatography on cobalt resin, followed by size exclusion chromatography and storage at −80 °C. The detailed methodology, which includes a TEV cleavage step to remove the His-tag, can be found elsewhere [68].

Primer and hairpin substrates (with 5′-FAM-label) and templates were purchased from Biomers.net. The duplexes were formed by mixing primer and template in a molar ratio of 1 to 1.5, in water containing 110 mM KCl. These mixtures were incubated for 10 min at 70 °C, then gradually cooled to 4 °C over a period of 2 h. The RdRp complex was obtained via stepwise mixing and incubation of the proteins [68], generating the nsp12-nsp7-nsp8 complex at 1:3:6 ratio and 10 μM concentration of nsp12. At the start of each assay, the RdRp complex was added to reaction buffer (= 20 mM HEPES pH 7.5, 50 mM NaCl and 5 mM MgCl_2_) and incubated with the appropriate RNA (sequence specified in each Figure panel) for 15 min at room temperature. To assess single nucleotide incorporation, we initiated the reaction by adding EDA-TP or ATP (at final concentrations given in the Figures) to reaction buffer and RdRp/RNA mix (final concentrations: 0.5 μM nsp12 and 0.5 μM RNA). When assessing multiple incorporations up to full extension, the other three NTPs (= CTP, UTP and GTP) were present at 10 µM. The samples were incubated at 30 °C and, at the indicated time points (0.5-60 min), aliquots were taken out to stop the reaction with two volumes of FBD (formamide, 10 mM EDTA and bromophenol blue). For the zero time point, RNA and RdRp were first added to the FBD stop solution, followed by the NTP/EDA-TP mix. Samples were heated for 10 min at 70 °C and cooled on ice before analysis by denaturing polyacrylamide gel electrophoresis [(PAGE); 20% acrylamide, 7M Urea] using 33 x 42 cm gels (C.B.S. Scientific) in TBE buffer at 45 W. The product bands were visualized using an Amersham Typhoon™ Biomolecular Imager (Cytiva) and analyzed with ImageQuant software.

To conduct the excision assay, we first performed 60 min extension of the hairpin RNA by the RdRp, in the presence of 20 µM EDA-TP, ATP or RDV-TP. The samples were heat-denatured for 10 min at 70 °C, after which the hairpin RNA was allowed to reanneal during 30 min incubation at 30 °C. Next, the ExoN complex [= nsp14 and nsp10 proteins combined in a molar ratio of 1:5 [68]] was added (final concentrations 50 nM nsp14 and 250 nM nsp10) and the reaction was incubated at 30 °C, with sample collection at 2 to 60 min. After stopping the reaction with FBD, gel electrophoresis was conducted as above.

### Primer extension in function of EDA-TP or NTP concentration

The pRSFDuet-1 plasmid encoding all three proteins (i.e. nsp7, nsp8 and nsp12) from SARS-CoV-2 Wuhan-Hu-1 was obtained from Addgene (ref.165451). Protein expression in *E. coli* and purification were performed as described [69]. The RdRp complex produced by this one-plasmid expression system is presumed to have the (nsp7)(nsp8)_2_(nsp12) stoichiometry [69] and still contains the 14His-tag at the N-terminus of nsp8.

Different primers (with 5′-FAM-label) and templates were purchased from Integrated DNA Technologies. The duplexes were formed by adding equimolar amounts of primer and template to annealing buffer, consisting of 10 mM Tris-HCl pH 7, 50 mM NaCl and 0.1 mM EDTA. These mixtures were heated for 3 min at 75 °C, then gradually cooled to 4 °C over a period of 2 h. (i) To measure the incorporation of a single EDA-TP or ATP, the primer:template was added to reaction buffer (100 mM NaCl, 20 mM HEPES pH 7.5, 10 mM MgCl_2_ and 5% glycerol), followed by the RdRp (final concentrations: 1.25 µM RNA and 2 µM enzyme). The reaction was initiated with 10 µM UTP and incubated for 10 min at 37 °C. Next, EDA-TP or ATP was added at different concentrations (range: 0.03-16 µM) and allowed to incorporate during 30 s. (ii) To measure CTP incorporation after EDA, we first allowed 10 min extension with 20 µM EDA-TP or ATP, to then add CTP (range: 0.06-100 µM). After (i) or (ii), the reaction was stopped with stop buffer (final concentration: 140 mM EDTA, 1.5 mM Tris-HCl, 49% formamide, 9% glycerol, 0.005% bromophenol blue and 0.005% xylene cyanol), followed by 10 min heating at 95 °C and immediate cooling on ice. After loading the samples on 13.3 × 8.7 cm pre-cast gels (Bio-Rad; 15% polyacrylamide and 7 M urea in TBE buffer), the gels were run in TBE buffer at 50 °C, for 30 min at 10 W. Visualization of the bands and analysis of the band intensities were done as described above.

### Cryo-EM methods

#### Sample preparation and data collection

Cryo-EM samples of the SARS-CoV-2 replication-transcription complex (RTC) were prepared as previously described [70]. Briefly, purified nsp12 at a concentration of 4.8 mg/ml was incubated with nsp7 and nsp8 at 4 °C for 3 h, at a molar ratio of 1:2:2 in a buffer containing 20 mM HEPES, pH 7.0, 100 mM NaCl, and 4 mM MgCl_2_. After purifying the mixture by Mono Q 5/50 ion-exchange chromatography (GE Healthcare), a stable nsp12-nsp7-nsp8 complex was obtained. The template and primer oligoribonucleotides (Takara; sequences provided in S3 Table) were annealed by heating the solution to 70 °C and gradually cooling it to 4 °C. The annealed RNA scaffold was incubated with the nsp12-nsp7-nsp8 complex for 30 min at 25 °C to form C-RTC. Next, the mixture was incubated with 8 mg/ml nsp13 and 5 mg/ml nsp9 for 30 min at 30 °C to assemble the E-RTC. Finally, the RTC was mixed with 50 μM EDA-TP to assemble the RTC1:EDA complex; or with 50 μM EDA-TP and 50 μM CTP to assemble the RTC2:EDA complex.

In total, 3 μl of protein-RNA complex at 3 mg/ml (added with 0.025% DDM) was applied onto a H_2_/O_2_ glow-discharged, 300-mesh Quantifoil R 1.2/1.3 grid (Micro Tools GmbH). The grid was blotted for 3.0 s with a blot force of 0 at 8 °C and 100% humidity, and plunge-frozen in liquid ethane using a Vitrobot (Thermo Fisher Scientific). Cryo-EM data were collected with a 300 keV Titan Krios electron microscope (Thermo Fisher Scientific) and a Falcon 4i direct electron detector (Thermo Fisher Scientific). Images were recorded at 130,000x magnification and calibrated at a super-resolution pixel size of 0.93 Å/pixel. The exposure time was set to 4 s with a total accumulated dose of 50 electrons per Å^2^. All images were automatically recorded using EPU and were collected with a defocus range from −2.0 μm to −1.0 μm.

### Data analysis

Each movie was motion-corrected and dose-weighted by MotionCor2. The contrast transfer function (CTF) was estimated by patch CTF in cryoSPARC v4.4. For the RTC1:EDA complex, 4161 micrographs were manually selected for further data processing. All subsequent analyses were performed in cryoSPARC except for those specifically mentioned. EMD-30504 was used to generate templates for template picker. 962,500 particles were selected and extracted with 220 pixels (bin2). After multiple rounds of 2D classification, 136,082 particles were selected and subjected to ab initio reconstruction jobs to generate initial models. Particles were further selected using these initial models as 3D reference for heterogeneous refinement resulting in 87,372 particles exhibiting RTC feature. Particles were finally re-extracted with 440 pixels and subjected to non-uniform refinement, yielding a reconstruction of RTC1:EDA with a resolution of 2.93 Å.

Using particle sorting workflow consistent with that described above, 106,477 particles were selected for NU refinement, resulting in an RTC2:EDA resolution of 2.78 Å. The local resolution of all final maps were estimated in cryoSPARC.

To build the structure of the SARS-CoV-2 RTC1:EDA and RTC2:EDA complexes, the coordinate of the SARS-CoV-2 RTC (PDB: 7CYQ) [70] was individually placed and fitted by rigid-body refinement into the cryo-EM map using UCSF Chimera. These models were manually built with EDA, using Coot with the guidance of the cryo-EM map, and in combination with real space refinement using Phenix. Statistics for data collection and refinement are provided in S5 Table.

### Data visualization and statistical analysis

Data were visualized and analyzed using GraphPad Prism v11. Dose-response curves were fitted by nonlinear regression to derive EC_50_, EC_99_, CC_50_, half-maximal and plateau values. Data are shown as mean ± SEM; individual data points represent independent experiments, with the number of experiments and statistical tests specified in the corresponding Figure legends. Statistical significance was defined as p ≤ 0.05. Where applicable, significance levels are indicated as *p ≤ 0.05, **p ≤ 0.01, ***p ≤ 0.001 and ****p ≤ 0.0001.

## Acknowledgments

We would like to thank Ashleigh Shannon for helpful discussions; Abhi Singh and Mizar Oliva for help with RdRp enzyme purification; Qingfeng Li for synthesizing EDA and its structural analogs; and Frédérick Coosemans for synthesizing the EDA-template. We thank the State Key Laboratory of Medicinal Chemical Biology at Nankai University for providing support on cryo-EM data collection.

## Data availability

All data needed to evaluate and reproduce the results in the paper are present in the paper and/or the Supplementary Materials. Coordinates and structure factors of the RTC1:EDA and RTC2:EDA have been deposited in the PDB with IDs 26HW and 26IA, respectively, and in EMDB with IDs 80666 and 80670, respectively.

