## Supplemental Figure 1 for "Mechanistic insights into coronavirus polymerase inhibition by a base-modified adenosine analog"

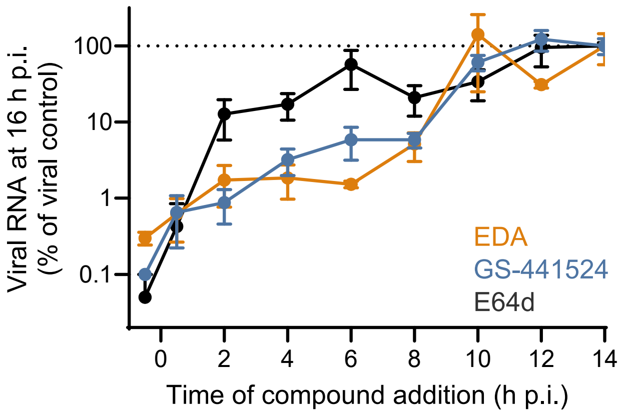


**S1 Fig. Time-of-compound addition assay.** HEL299 cells were infected with HCoV-229E and the compounds were added to separate wells at -0.5, 0.5, 2, 4, 6, 8, 10, 12 or 14 h p.i. At 16 h p.i., the cells were washed with ice-cold PBS, then mixed with Cells-to-cDNA II lysis buffer and incubated for 10 min at 75 °C. The viral RNA copy number was determined by one-step RT-qPCR as explained in the main methodology section. The input RNA (= starting number of viral RNA copies) was derived from virus-infected cells that received no compound and underwent lysis at 2 h p.i. Data shown are the mean ± SEM (N=4).
