## Supplemental Figure 2 for "Mechanistic insights into coronavirus polymerase inhibition by a base-modified adenosine analog"

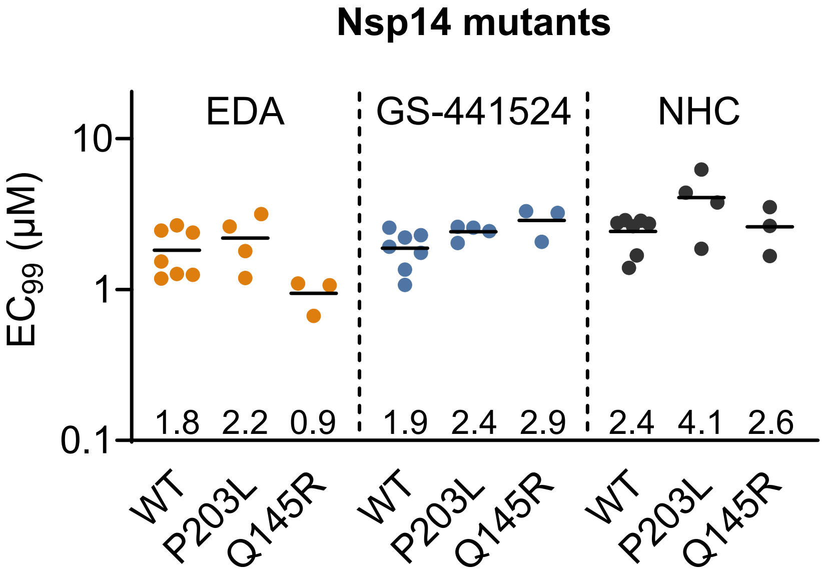


**S2 Fig. Evaluation against engineered SARS-CoV-2 viruses bearing mutations in nsp14.** The Y-axis shows $EC_{99}$ values determined by RT-qPCR for viral load. Dot symbols represent individual data points from 3-7 independent experiments. Mean values are shown as bars and are also indicated above the X-axis. Mutants were compared with WT by ordinary one-way ANOVA with Dunnett's multiple-comparisons correction; no significant differences were observed.
