## Supplemental Figure 3 for "Mechanistic insights into coronavirus polymerase inhibition by a base-modified adenosine analog"

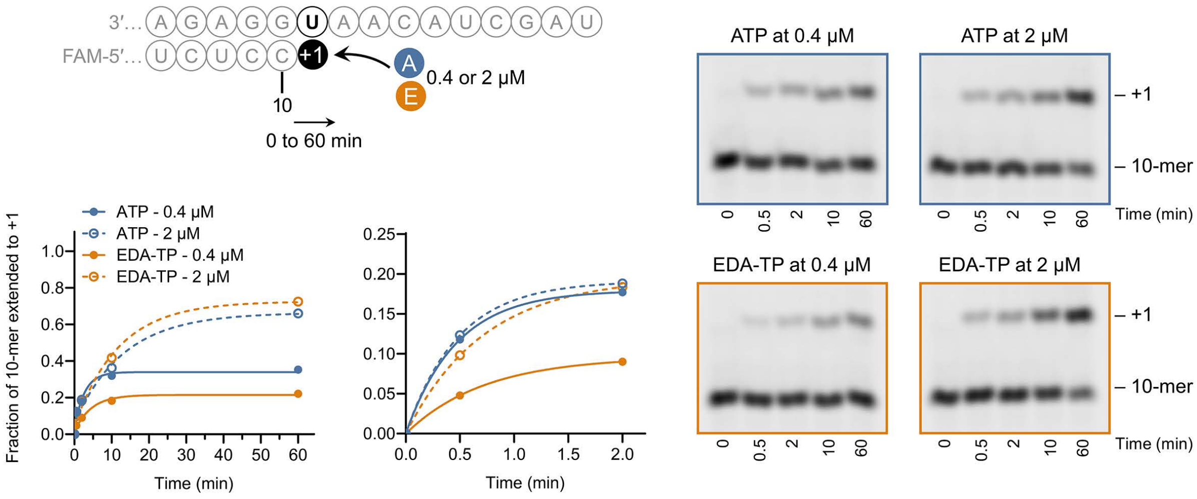


**S3 Fig. Time course for single-nucleotide incorporation of EDA-TP or ATP.** The graphs show the fraction of the primer extended with EDA-TP or ATP, within the full time frame of 0-60 min (left) or a short time frame of 0-2 min (right), to better visualize early product formation. Note that the Y-axis is different in the two graphs. Data points are the mean from two experiments.
