## Supplemental Figure 4 for "Mechanistic insights into coronavirus polymerase inhibition by a base-modified adenosine analog"

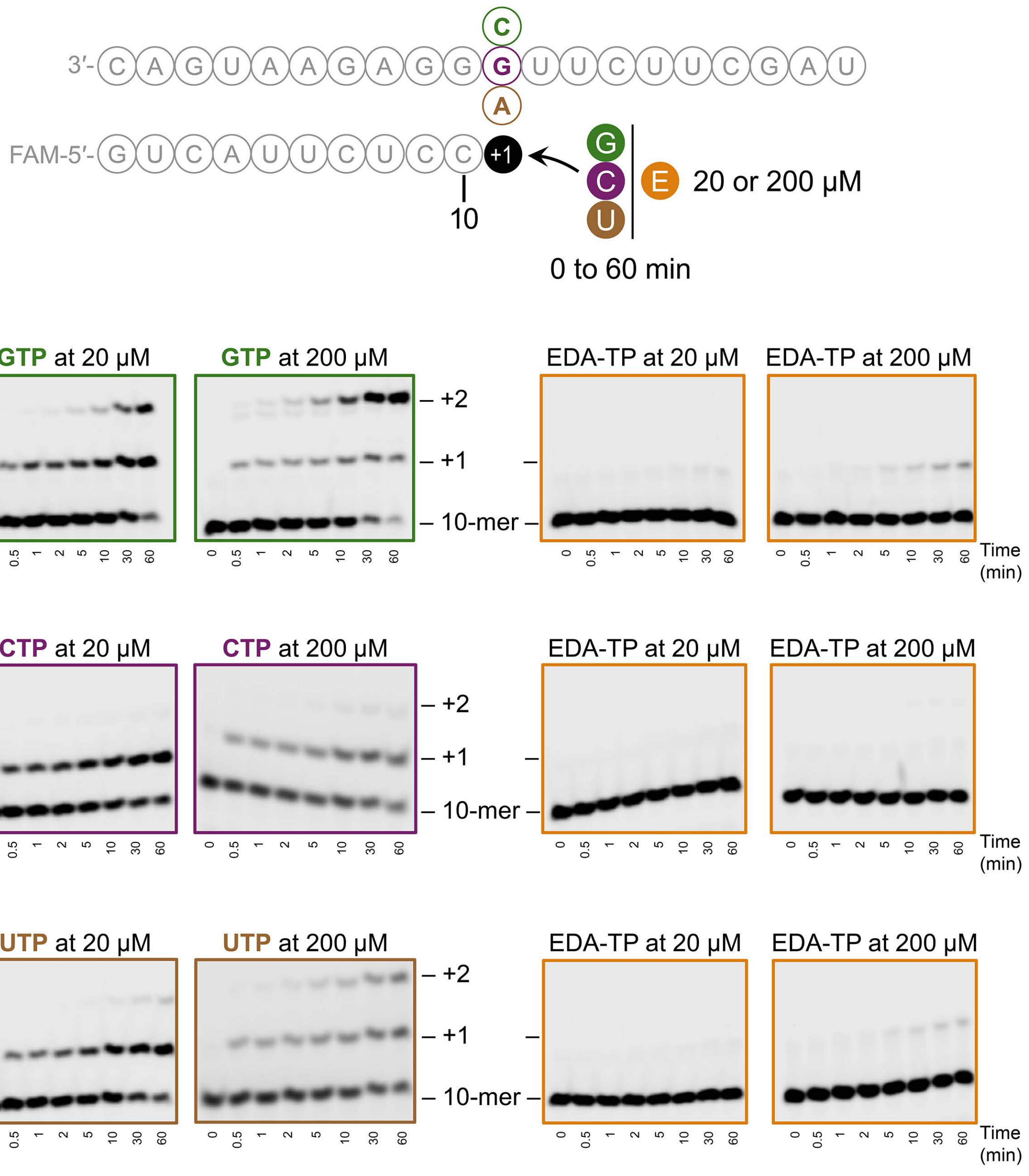


**S4 Fig. EDA-TP does not incorporate as a substitute for GTP, CTP nor UTP.** Time course for single-nucleotide incorporation of EDA-TP, tested at 20 or 200 µM, as a substitute for GTP (top panels; template with C); for CTP (middle panels; template with G); or for UTP (bottom panels; template with A). For the natural GTP, CTP and UTP substrates, the expected incorporation at the +1 position is followed by some level of mis-incorporation at the +2 position. This is clearest with GTP; an NTP concentration of 200 µM; and at longer reaction times. In sharp contrast, EDA-TP generates only very faint bands at +1, when tested at 200 µM and as a mimic of GTP or UTP.
