## Supplemental Figure 5 for "Mechanistic insights into coronavirus polymerase inhibition by a base-modified adenosine analog"

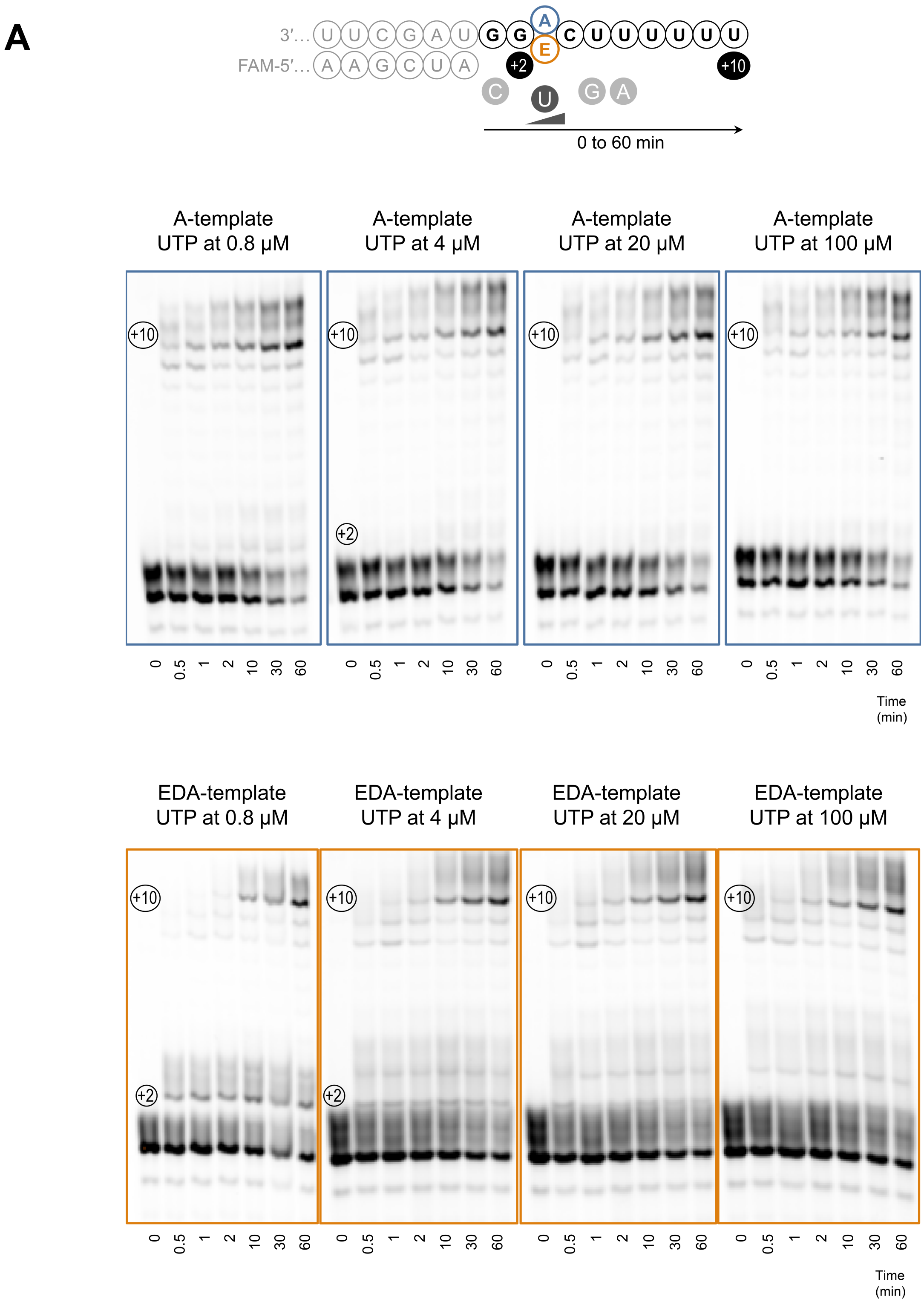
A
3′… U U C G A U G G A C U U U U U U
FAM-5′… A A G C U A E
+2
+10
C U G A
0 to 60 min
A-template UTP at 0.8 μM
A-template UTP at 4 μM
A-template UTP at 20 μM
A-template UTP at 100 μM
+10
+2
0 0.5 1 2 10 30 60
Time (min)
EDA-template UTP at 0.8 μM
EDA-template UTP at 4 μM
EDA-template UTP at 20 μM
EDA-template UTP at 100 μM

B

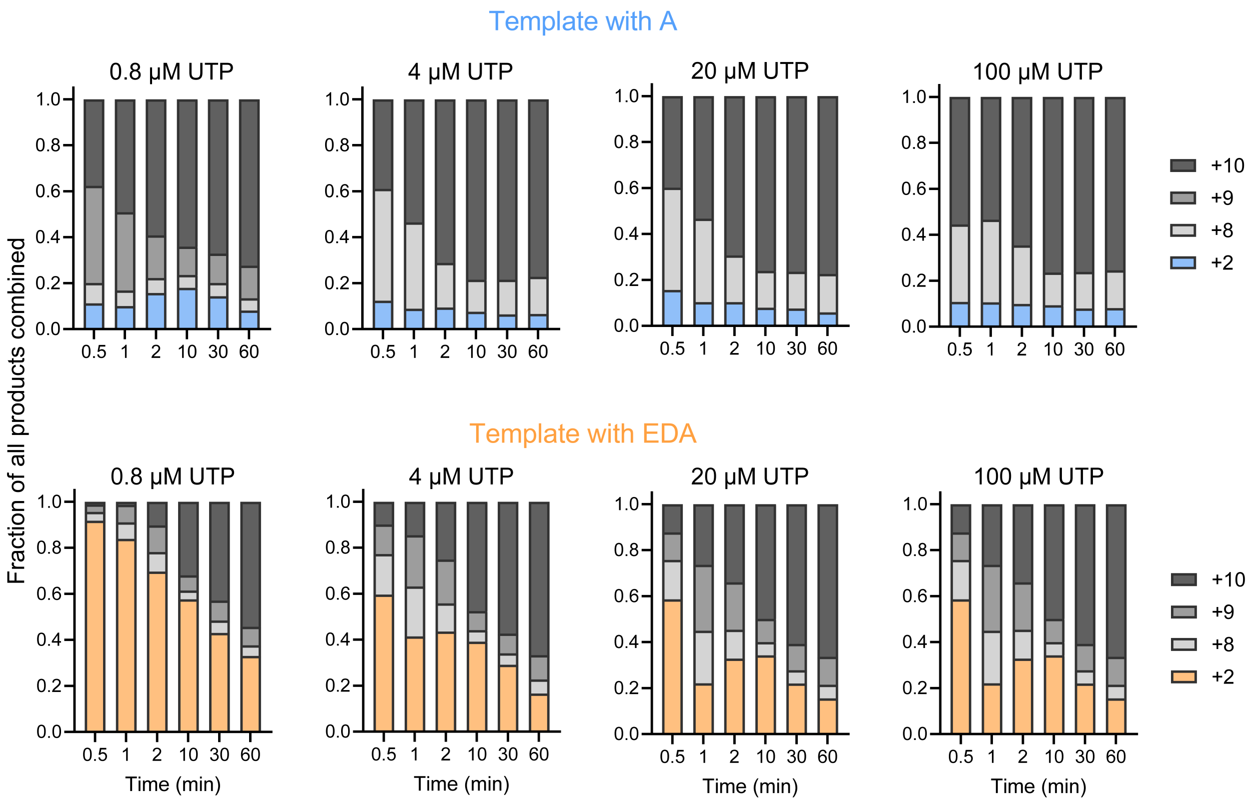


**S5 Fig. EDA stalls RNA synthesis when present in the template, but the effect is transient and dependent on the NTP concentration.** (**A**) Time-dependent polymerization against a template containing an A residue (gel images boxed in blue) or EDA residue (gel images boxed in orange), in the presence of 0.8, 4, 20 or 100 µM UTP. (**B**) The graph shows the relative band intensities of the +2 intermediate with two incorporated C residues, preceding addition of UTP against the template with A (blue) or the template with EDA (orange). The longer products are shown in different shades of grey. Data are from a single experiment.
