## Supplemental Table 1 for "Mechanistic insights into coronavirus polymerase inhibition by a base-modified adenosine analog"

**S1 Table. Evaluation of 6-alkyl and 6-aryl substituted 7-deazapurine ribonucleoside analogs in HCoV-229E-infected HEL299 cells.**

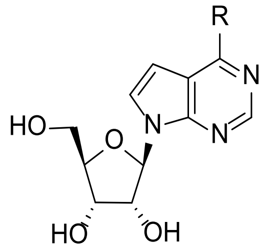


| Compound | R | **HCoV-229E** in HEL299 cells[a] | | |
|---|---|---|---|---|
| | | $EC_{50}$ (µM) | $CC_{50}$ (µM) | SI |
| EDA | 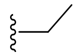 | 1.8 ± 0.1[b] | >100 | >56 |
| MeDA | 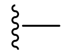 | 0.036 ± 0.002[b] | 0.64 ± 0.03[b] | 18 |
| **1** | 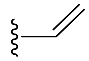 | >100 | >100 | - |
| **2** | 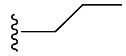 | 53 ± 4[b] | >100 | >2 |
| **3** | 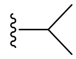 | 47 ± 7[b] | >100 | >2 |
| **4** | 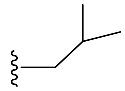 | >100 | >100 | - |
| **5** | 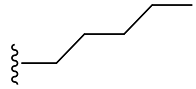 | >100 | >100 | - |
| **6** | 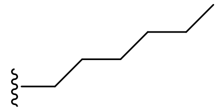 | >100 | 81 | - |
| **7** | 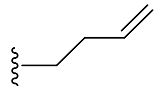 | >100 | >100 | - |
| **8** | 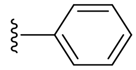 | >100 | >100 | - |
| **9** | 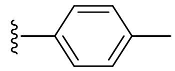 | >100 | >100 | - |
| **10** | 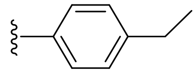 | 35 ± 4[b] | >100 | >3 |

| | | | | |
|---|---|---|---|---|
| **11** | 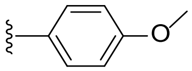 | >100 | >100 | - |
| **12** | 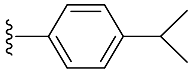 | >100 | 93 | - |
| **13** | 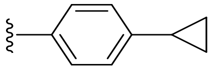 | >100 | >100 | - |
| **14** | 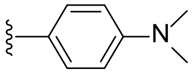 | >100 | >100 | - |
| **15** | 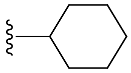 | 26 ± 3[b] | >100 | >4 |
| **16** | 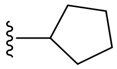 | >100 | >100 | - |
| Tubercidin | $-NH_2$ | 0.29 ± 0.03[b] | 2.0 ± 0.2[b] | 7 |
| GS-441524 | | 2.5 ± 0.1[b] | >100 | >40 |
| NHC | | 0.52 ± 0.13[b] | >100 | >192 |

[a]Unless specified otherwise, each analog was tested once.
[b]Mean value ± SEM (N=3-5).
$EC_{50}$: half-maximal effective concentration, determined by MTS-based CPE reduction assay.
$CC_{50}$: 50% cytotoxic concentration, determined in mock-infected cells using MTS cell viability assay.
SI: selectivity index, i.e. ratio of $CC_{50}$ over $EC_{50}$.
