## Supplemental Table 2 for "Mechanistic insights into coronavirus polymerase inhibition by a base-modified adenosine analog"

**S2 Table. Anti-CoV evaluation of analogs bearing a short alkyl group with various heteroatom substitutions.**

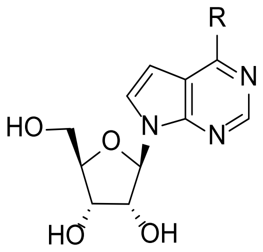

| Compound | R | HCoV-229E in HEL299 cells[a] | | | SARS-CoV-2 in A549[ACE2/TMPRSS2] cells[a] | | |
|---|---|---|---|---|---|---|---|
| | | $EC_{50}$ (µM) | $CC_{50}$ (µM) | SI | $EC_{50}$ (µM) | $CC_{50}$ (µM) | SI |
| **17** | 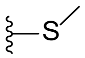 | >100 | >100 | - | 66 ± 66[b] | >100 | >2 |
| **18** | 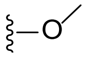 | >100 | >100 | - | 50 ± 10[b] | >100 | >2 |
| **19** | 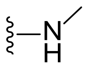 | >100 | >100 | - | >100 | >100 | - |
| **20** | 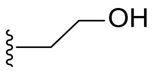 | 15 ± 1[b] | >100 | >7 | 15 ± 2[b] | 72 ± 8[b] | 5 |
| **21** | 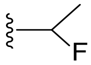 | >100 | >100 | - | 17 ± 5[b] | >100 | >6 |
| **22** | 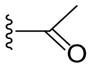 | >100 | >100 | - | >100 | >100 | - |
| **23** | 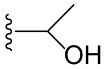 | >100 | >100 | - | >100 | >100 | - |
| **24** | 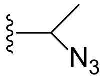 | >100 | >100 | - | 76 ± 22[b] | >100 | >1 |
| **25** | 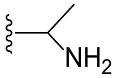 | >100 | >100 | - | >100 | >100 | - |

[a]Unless specified otherwise, each analog was tested twice.
[b]Mean value ± SEM (N=3).
$EC_{50}$: half-maximal effective concentration, determined by MTS-based CPE reduction assay.
$CC_{50}$: 50% cytotoxic concentration, determined in mock-infected cells using MTS cell viability assay.
SI: selectivity index, i.e. ratio of $CC_{50}$ over $EC_{50}$.
