## Supplemental Table 3 for "Mechanistic insights into coronavirus polymerase inhibition by a base-modified adenosine analog"

**S3 Table. Sequences of the primer:template and hairpin oligonucleotides used in the enzymatic and cryo-EM studies.**

| Purpose | Sequence |
|---|---|
| **Primer-extension assay** | |
| Fig. 5A | 3′-CAGUAAGAGGAUUCUUCGAUAAUGCUAGUGUACCCCUAUC-5′<br>FAM-5′-GUCAUUCUCCUAAGAAGCUA-3′ |
| Fig. 5B | FAM-5′-CGAGAGAC**U**CGCGUAGUUUUCUACGCG-3′ (hairpin with U)<br>FAM-5′-CGAGAGA**UU**CGCGUAGUUUUCUACGCG-3′ (hairpin with UU) |
| Fig. 5C | FAM-5′-CGAGAGAGUCGCGUAGUUUUCUACGCG-3′ (hairpin with U) |
| Fig. 6A | 3′-CAGUAAGAGGAUUCUUCGAUAAUGCUAGUGUACCCCUAUC-5′<br>FAM-5′-GUCAUUCUCCUAAGAAGCUA-3′ |
| Fig. 6B | 3′-CAGUAAGAGGUAACAUCGAU-5′<br>FAM-5′-GUCAUUCUCC-3′ |
| Fig. 6C | 3′-CAGUAAGGGAUUCUUCGAUGG**A**CUUUUUU-5′ (template with A)<br>FAM-5′-GUCAUUCCCUAAGAAGCUA-3′<br><br>3′-CAGUAAGGGAUUCUUCGAUGG**E**CUUUUUU-5′ (template with EDA)<br>FAM-5′-GUCAUUCCCUAAGAAGCUA-3′ |
| Fig. S3 | 3′-CAGUAAGAGGUAACAUCGAU-5′<br>FAM-5′-GUCAUUCUCC-3′ |
| Fig. S4 | 3′-CAGUAAGAGG**C**UUCUUCGAUU-5′<br>FAM-5′-GUCAUUCUCC-3<br><br>3′-CAGUAAGAGG**G**UUCUUCGAU-5′<br>FAM-5′-GUCAUUCUCC-3′<br><br>3′-CAGUAAGAGG**A**UUCUUCGAU-5′<br>FAM-5′-GUCAUUCUCC-3′ |
| Fig. S5 | 3′-CAGUAAGGGAUUCUUCGAUGG**A**CUUUUUU-5′ (template with A)<br>FAM-5′-GUCAUUCCCUAAGAAGCUA-3′<br><br>3′-CAGUAAGGGAUUCUUCGAUGG**E**CUUUUUU-5′ (template with EDA)<br>FAM-5′-GUCAUUCCCUAAGAAGCUA-3′ |
| **Cryo-EM analysis** | |
| RTC1:EDA | 3′-UAAUUUCCAAAUAUGGAAGGGUCCAUUGUUUGGUUGGA**UUU**AAACUAGAGAACAUCUAG-5′<br>5′-AUUAAAGGUUUAUACCUUCCCAGGUAACAAACCAACCU-3′ |
| RTC2:EDA | 3′-UAAUUUCCAAAUAUGGAAGGGUCCAUUGUUUGGUUGGA**UGC**AAACUAGAGAACAUCUAG-5′<br>5′-AUUAAAGGUUUAUACCUUCCCAGGUAACAAACCAACCU-3′ |
