## Supplemental Table 4 for "Mechanistic insights into coronavirus polymerase inhibition by a base-modified adenosine analog"

**S4 Table. Primers and probes.**

| | |
|---|---|
| **HCoV-229E** | |
| Primer 1 | 5′-TTAGAGAGCGTGTTGAAGGTG-3′ |
| Primer 2 | 5′-GTTCTGAATTCTTGCGCCTAAC-3′ |
| Probe | 5′-[FAM]-TCTGGGTTG-[ZEN]-CTGTTGATGGTGCTA-[IABkFQ]-3′ |
| **HCoV-OC43** | |
| Primer 1 | 5′-CCAAGTAGCGATGAGGCTATTC-3′ |
| Primer 2 | 5′-GCGCGAAGTAGATCTGGAATTA-3′ |
| Probe | 5′-[FAM]-ATTGAAGGC-[ZEN]-TGAGGAAGGTCTGCT-[IABkFQ]-3′ |
| **MHV-A59** | |
| Primer 1 | 5′-GGAGACAGCATTGAAGGAGTC-3′ |
| Primer 2 | 5′-CTTGGGTCCCTTTCGACAATA-3′ |
| Probe | 5′-[FAM]-AACAGCCAA-[ZEN]-GCGGACACCAATACC-[IABkFQ]-3′ |
| **SARS-CoV-2** | [1] |
| *20A.EU2* | |
| Primer 1 | 5′-GACCCCAAAATCAGCGAAAT-3′ |
| Primer 2 | 5′-TCTGGTTACTGCCAGTTGAATCTG-3′ |
| Probe | 5′-[FAM]-ACCCCGCAT-[ZEN]-TACGTTTGGTGGACC-[IABkFQ]-3′ |
| *Omicron* | |
| Primer 1 | 5′-TTACAAACATTGGCCGCAAA-3′ |
| Primer 2 | 5′-GCGCGACATTCCGAAGAA-3′ |
| Probe | 5′-[FAM]-ACAATTTGC-[ZEN]-CCCCAGCGCTTCAG-[IABkFQ]-3′ |

**1.** Vogels CBF, Brito AF, Wyllie AL, Fauver JR, Ott IM, Kalinich CC, et al. Analytical sensitivity and efficiency comparisons of SARS-CoV-2 rt-qpcr primer-probe sets. Nat Microbiol. 2020;5(10):1299–305. doi: 10.1038/s41564-020-0761-6.
