## Supplemental Table 5 for "Mechanistic insights into coronavirus polymerase inhibition by a base-modified adenosine analog"

**S5 Table. Cryo-EM data collection, refinement and validation statistics.**

| | RTC1:EDA | RTC2:EDA |
|---|---|---|
| PDB entry | 26HW | 26IA |
| EMDB entry | 80666 | 80670 |
| **Data collection and processing** | | |
| Magnification | 130,000 | 130,000 |
| Voltage (kV) | 300 | 300 |
| Electron exposure ($e^{-}/Å^2$) | 50 | 50 |
| Defocus range (μm) | -1.0 to -2.0 | -1.0 to -2.0 |
| Pixel size (Å) | 0.93 | 0.93 |
| Symmetry imposed | C1 | C1 |
| Initial particle images (no.) | 136,082 | 154,208 |
| Final particle images (no.) | 87,372 | 106,477 |
| Map resolution (Å) | 2.93 | 2.78 |
| FSC threshold | 0.143 | 0.143 |
| Map resolution range (Å) | 2.93 – ∞ | 2.78 – ∞ |
| **Refinement** | | |
| Model composition | | |
| Non-hydrogen atoms | 21,485 | 21,488 |
| Protein residues | 2,654 | 2,654 |
| Ligands | 1 | 1 |
| *B* factors ($Å^2$) | | |
| Protein | 36.40 | 63.41 |
| Ligand | 74.27 | 102.49 |
| R.m.s. deviations | | |
| Bond lengths (Å) | 0.006 | 0.010 |
| Bond angles (°) | 0.702 | 0.707 |
| Validation | | |
| MolProbity score | 2.70 | 2.81 |
| Clashscore | 54.45 | 52.76 |
| Poor rotamers (%) | 1.98 | 2.47 |
| Ramachandran plot | | |
| Favored (%) | 96.12 | 95.44 |
| Allowed (%) | 3.80 | 4.48 |
| Disallowed (%) | 0.08 | 0.08 |
